# Hydraulic Activation of the AsLOV2 Photoreceptor

**DOI:** 10.1101/2025.06.19.660617

**Authors:** Shiny Maity, Jackson Sheppard, Hannah Russell, Chung-Ta Han, Leah Epstein, Bruce A. Johnson, Brad Price, Ruixian Han, Lokeswara Rao Potnuru, Jinlei Cui, Mark S. Sherwin, Joan-Emma Shea, Janet E. Lovett, Kevin H. Gardner, Songi Han

## Abstract

How proteins transduce light into mechanical energy remains a central question in biology. This study tests the hypothesis that blue light activation of the LOV2 (light, oxygen, voltage sensitive) domain of *Avena sativa* phototropin 1 (AsLOV2), gives rise to concerted water movement that induces protein conformational extensions. Using electron and nuclear magnetic resonance spectroscopy, along with molecular dynamics simulations at high pressure, we find AsLOV2 activation can be initiated by blue light or high pressure, followed by selective and concerted expulsion of low-entropy, tetrahedrally coordinated “wrap” water from the protein hydration shell. These findings suggest that interfacial water serves as constituents to reshape the protein’s free energy landscape during activation. Our study highlights hydration water as an active hydraulic fluid that can drive long-range conformational changes underlying protein mechanics upon light activation and offers a new concept for engineering externally controllable protein actuators.

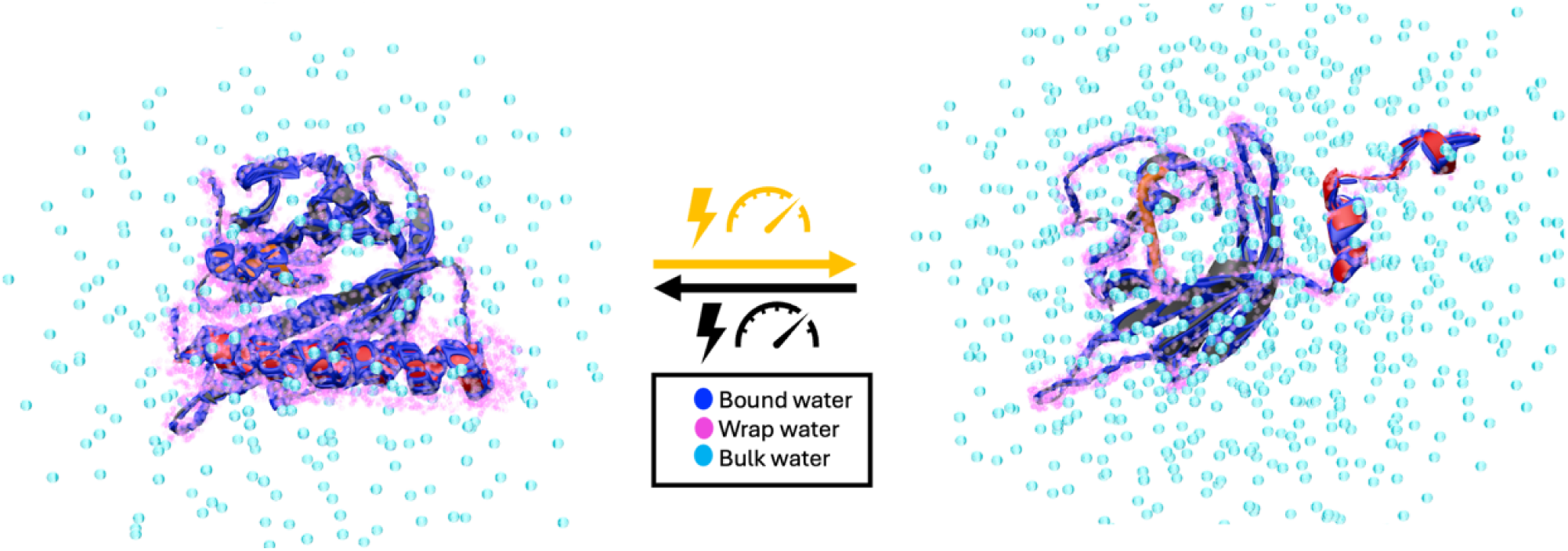

## Introduction

Flavin-binding photoreceptors are critical components across bacteria, plants, animals, and humans, facilitating essential signal transduction pathways and governing vital biological processes such as circadian rhythms and phototropism(*1–4*). Among flavin-containing photoreceptors, the Light, Oxygen, or Voltage (LOV) sensing flavoproteins are particularly notable for their mechanical response to blue-light stimulation, initiating cascades of biochemical and mechanical signaling events through structural rearrangements. The ability of LOV proteins to convert absorbed photon energy into controlled mechanical actuation of an associated kinase makes them some of the most attractive protein actuator candidates for bioengineering applications. They have been engineered to control cellular signaling because of their remarkable ability to “turn on” the activity of a tethered protein, simply by shining light. Today, engineered LOV variants are widely used to regulate diverse cellular processes—including cell motility (PA RAC1)(*5*), nuclear transport (LINuS/LEXY)(*6*, *7*), gene expression (CLASP)(*8*), ion flux (BLINK)(*9*), proteolytic cleavage (FLARE)(*10*), organelle transport (TULIP)(*11*), prokaryotic transcription (LOVTAP)(*12*), and developmental signaling (ilid SOS)(*13*). The LOV2 domain of *Avena sativa* phototropin 1 (AsLOV2) exemplifies this capability that utilizes an oxidized flavin mononucleotide (FMN) chromophore non-covalently bound within its structure. Upon illumination with blue light (λ = 450 nm), the FMN cofactor transitions to a triplet state and forms a covalent bond with a conserved cysteine residue (C450) that reduces the flavin isoalloxazine ring, triggering conformational changes in AsLOV2 that mechanically actuate a tethered effector protein, thereby optically driving the effector protein function and correlated cellular processes(*14–17*).

Light activation offers substantial advantages compared to chemical modulation, among others, due to its superior temporal and spatial resolution(*6*, *8*, *13*). This method enables precise and rapid initiation and termination of biological responses, which are inherently reversible and repeatable without cumulative chemical side-effects or undesirable off-target interactions. For AsLOV2-derived tools, much of this regulation starts with light-driven protein conformational changes triggered by the photochemical events at the chromophore. While high-resolution X-ray crystallography studies of illuminated AsLOV2 do not show any long-range displacement from the dark state structure(*18*), multiple spectroscopic approaches including high-resolution nuclear magnetic resonance (NMR), time-resolved electron paramagnetic resonance (EPR), and Fourier-transform infrared (FTIR) spectroscopy demonstrated that light activation of LOV is accompanied by partial unfolding of the A’α and Jα helix and displacement of the Jα helix from the Iβ-sheet surface, suggesting a plausible mechanism for transmitting structural changes(*19–24*).For LOV domains to function as actuators, mechanical force must arise from coordinated conformational changes that propagate from the chromophore to the A’α and Jα helices then to downstream signaling partners. Yet, the molecular events orchestrating such concerted motions remain unidentified. Water has long been implicated in LOV activation for acting as an intrinsic base catalyzing redox reactions in the context of the FMN photocycle (*25–30*). However, the role of water as a mechanical modulator from concerted movement, beyond that of a proton donor/acceptor, has not been strongly considered. Rather than acting solely as a chemical cofactor, water may participate directly in facilitating long-range conformational changes that underpin mechanical signaling. A series of studies reported on collective water movement driving activation in different protein systems, including tryptophan excitation followed by femtosecond spectroscopy(*31–34*), enzyme activation by water-mediated thermal network(*32*), and water network mediated protein dynamical transitions(*35*), but these are still emerging concepts which remain under refinement. Direct studies of both protein and water movement upon protein activation are needed to obtain further clarity, both in general and in this particularly useful photoreceptor.

This study tests our hypothesis that collective action of water hydrating AsLOV2, initiated by a thermodynamic state change, drives structural transitions of the LOV domain. We investigate changes in the hydration water landscape, concurrent with out-of-equilibrium conformational changes of AsLOV2 initiated by blue-light activation, using advanced spectroscopic and computational methodologies. To monitor light-induced conformational changes, we used time-resolved EPR for real-time protein motion analysis and double electron-electron resonance (DEER) spectroscopy for quantifying intra-protein distance changes. To assess light-induced changes in near-protein water, we employed ^17^O NMR chemical shift analysis to resolve distinct hydration water populations and Overhauser dynamic nuclear polarization (ODNP) to probe diffusion dynamics of water at sub-nanosecond timescales. To evaluate pressure-induced effects, we performed high-pressure DEER to monitor conformational extension, high-pressure solution-state NMR to detect residue-specific structural responses, and atomistic molecular dynamics (MD) simulations to model pressure-driven changes in protein conformation and hydration. Together, these measurements reveal a new perspective into concurrent molecular level changes of the protein and water constituents of AsLOV2 upon light and pressure activation.

We look to identify distinct populations of hydration water at the protein-water interface, including water tightly bound to the protein surface and lower density, lower entropy, tetrahedrally structured, “wrap” water hydrating small hydrophobic regions of AsLOV2(*36–43*), and uncover the consequences of light- and/or pressure-induced changes in the population and dynamics of protein-adjacent water and protein conformations. This study tests a new hypothesis that AsLOV2 operates as a hydraulic actuator in which collective movement of hydration water induces protein conformational transitions. This study compels us to consider a novel conceptual framework for light-driven mechanical actuation of photosensory domains in the LOV class as used in plant phototropins as well as numerous fungal and bacterial proteins(*44–47*).

## Results

### Significant conformational extension in WT and N414Q AsLOV2 upon light activation

To elucidate the molecular mechanism underlying the light-induced mechanical action of AsLOV2, we examined the wild-type (WT) protein and the slow-cycling N414Q variant. The N414 residue was thought to be compulsory in the AsLOV2 activation process(*18*, *23*, *28*, *48*, *49*), but the discovery that LOV domains lacking N414 still show strong activity, except with slower photocycle times (Vvd: 18,000 s; YtvA: 3,600 s), gave rise to the hypothesis that water may regulate the N5 proton transfer in FMN as a proton acceptor, with N414 involved remotely from over 10 Å away (*28*, *46*, *48*). We selected the N414Q mutant to compare to WT to examine the role of water in mediating AsLOV2 function.

Time-resolved UV-Vis measurements confirmed that the FMN dark-state recovery kinetics in the N414Q variant was slowed by a factor of 2.5 (***τ***FMN_N414Q = 172.80 ± 0.02 s) compared to WT AsLOV2 (***τ***FMN_WT = 67.97 ± 0.03 s), while the overall amplitude change in UV-Vis absorption is comparable between the variants, consistent with the literature(*28*) (Fig. 1A, B). To investigate the mechanical response of AsLOV2 upon photoactivation in real-time under physiological conditions, we employed time-resolved gadolinium-gadolinium electron paramagnetic resonance (TiGGER) spectroscopy at 240 GHz(*21*, *50*). The results confirmed that both variants exhibit significant and comparable conformational extensions upon blue light illumination, as observed by changes in the dipolar coupling between Gd-TPATCN spin-labels introduced at residues T406C and E537C, located on the N-terminal A’α and C-terminal Jα helices, respectively (SI Fig. S3).

**Figure 1.**
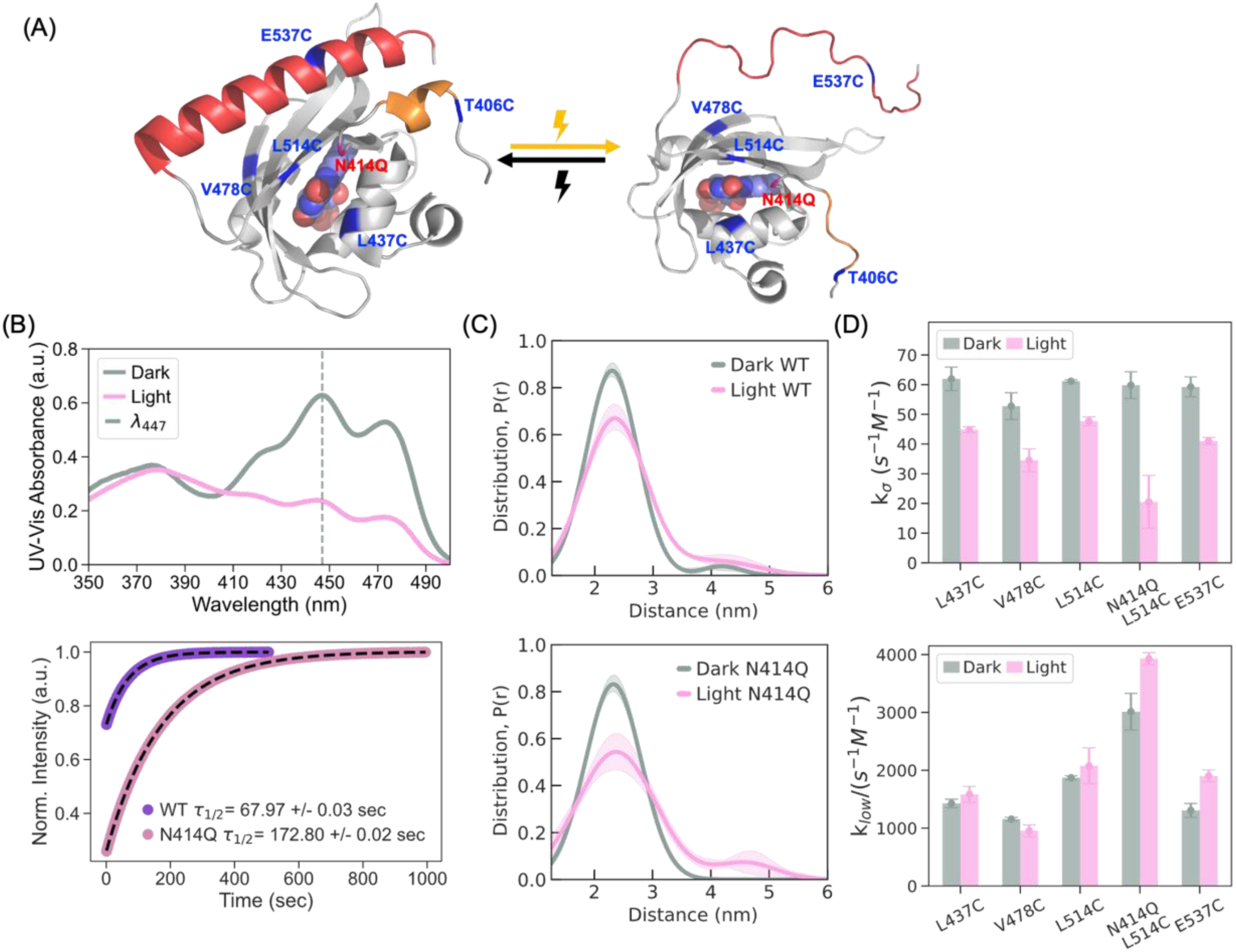
Photoactivation of AsLOV2 induces long range conformational change and modulates interfacial hydration dynamics. (A) Crystal structure of AsLOV2 in the dark state (PDB: 2V1A(*62*)), highlighting sites labeled with MTSL for ODNP, TiGGER, and DEER measurements (left). Upon photoactivation, the A’α (orange) and Jα helices (red) unfold and move further apart, as illustrated on the right. (B) UV-Vis absorbance spectroscopy confirms light-induced activation of AsLOV2. The wild-type protein exhibits a decrease in absorbance at ~450 nm under blue light illumination (top). In the N414Q mutant, time-resolved measurements at 450 nm reveal a 2.5-fold slower return to the dark state, indicating a stabilized lit-state conformation (bottom). (C) The distance distributions obtained with a DeerLab two-Gaussian model (see SI for further details) from the DEER measurements between T406C and E537C show an increase in the population of extended conformations (3–5 nm) in the lit state (top). The N414Q mutant further enriches the extended state population under illumination, consistent with enhanced structural stabilization of the lit conformation (bottom). (D) ODNP measurements reveal that light activation leads to a strong decrease in fast-moving water (k_σ_) across all sites, and a smaller increase in slow-moving water (*k*_low_) across most sites, suggesting reduced hydration dynamics at the protein interface. This effect is accentuated in the N414Q mutant, particularly at site L514C, where further reduction in k_σ_ and increase in *k*_low_ indicate enhanced local dehydration and stronger stabilization of the photoactivated state.

To evaluate the extent of distance changes, we performed DEER(*51–54*) at cryogenic temperatures on both variants that are spin-labeled with MTSL (*S*-(1-oxyl-2,2,5,5-tetramethyl-2,5-dihydro-1H-pyrrol-3-yl) methyl methanesulfonothioate) at T406C and E537C. Samples were photoactivated in solution at room temperature prior to freezing to accumulate lit-state populations and to preserve the physiologically relevant lit-state structure. DEER of the dark state showed a mean distance at 2.2 nm between the spin labels at sites 406 and 537 in both variants. Upon photoactivation, the amplitude at 2.2 nm dropped with a distance shift toward 3–5 nm to comparable extent in the WT and N414Q variant, indicating light-induced conformational extension in both proteins. (Fig. 1A, C). The observed structural rearrangements in solution state encompassing 1-3 nanometer-scale displacements contrast with X-ray crystal structures that only show sub-angstrom displacements between T406C and E537C but are consistent with other solution spectroscopic results. To fully resolve the structural transitions and associated hydration changes upon AsLOV2 activation, we require structural biology tools capable of probing out-of-equilibrium states in solution.

### Reduced surface hydration water dynamics upon light activation

To examine changes in the property of hydration water upon light activation in solution state, we relied on ODNP using MTSL at selected AsLOV2 sites to experimentally evaluate the local diffusion dynamics of hydration water associated with AsLOV2. ODNP probes site-specific water dynamics within ~1 nm of the spin label, capturing diffusion processes with correlation times on the order of tens of picoseconds to sub-nanoseconds(*55–57*). We examined four sites: E537C on the Jα helix, which unfolds upon photoexcitation; L514C on the Iβ sheet, located near the conserved glutamine Q513; L437C on the Cα helix; and V478C on the Gβ sheet (Fig. 1A).

ODNP leverages dipolar interaction between the unpaired electron spin (e) of MTSL and the ^1^H nuclei of surrounding water molecules. This interaction results in preferential e-^1^H cross relaxation via the double quantum transition (*w_2_*) that is facilitated when the instantaneous motion of water near the spin label occurs on a timescale comparable to or faster than the inverse electron spin Larmor frequency of 9.8 GHz at 0.35 T. Efficient e-^1^H *w_2_* transition followed by physical diffusion of the polarized water from near the spin label to bulk water results in ^1^H NMR signal enhancement of the bulk water signal owing to greater thermal polarization of the electron spin by 658 fold than the ^1^H nuclear spin. By analyzing ODNP enhancements (ε(p)) and ^1^H NMR *T*_1_ relaxation times, we can extract the coupling factor (ξ) that yields the e-^1^H dipolar correlation times and a local surface water diffusivity, *D*_local_. Further analysis of the ODNP data allows us to extract two key parameters that report on surface water properties at very different time scales: the cross-relaxivity parameter (*k****_σ_***) reflecting on water motion across the dipolar field of the spin label on the picosecond timescale, dictated by the EPR frequency of the spin label (9.8 GHz), and the self-relaxivity parameter (*k*_low_) capturing bound water population moving on the tens of nanoseconds timescale, primarily influenced by the fluctuation of interactions at the NMR frequency (14.8 MHz)(*55–60*).

The EPR lineshape of the spin label at site E537C on the Jα helix shows motional narrowing in the lit-compared to dark state (SI Fig.S1), suggesting increased side chain dynamics. This is consistent with insight from the literature that the Jα helix unfolds upon light activation(*19*, *21*, *22*, *49*, *61*). Based on that, site E537C should experience increased exposure to freely exchanging water upon unfolding. Spin labels introduced at residues L437C, V478C, and L514C, all of which are on the same face of AsLOV2 as the Jα helix, showed minimal change in the EPR lineshape upon light activation. Across all sites examined, we observed reduced diffusion dynamics of water near the protein surface: *D*_local_ decreased around all sites (SI Fig. S2), accompanied by reduced *k****_σ_*** (Fig. 1D top), indicating a slowdown in surface water diffusivity at the protein interface and an increased *k*_low_ for site E537C (Fig. 1D bottom) indicating a greater population of slow moving, bound, water population upon light activation at these sites. Similar trends were observed in the slow-cycling N414Q mutant at site L514C, with greater changes measured in its hydration environment compared to WT upon light activation (Fig. 1D). For hydration water dynamics to slow down at site E537C that is experiencing less restriction in movement and greater accessibility to water, the composition of hydration water around this site must be altered towards an increased fraction of hydration water population with slower mobility upon light activation.

We hypothesize that at least two distinct water populations besides bulk water make up the hydration layer of AsLOV2, (i) hydrogen-bonded (bound) water with slower dynamics and (ii) water with greater tetrahedral structuring and lower entropy wrapping over small hydrophobic protein sites (wrap water)—a categorization consistent with recent studies reporting on high-density, icosahedral and low-density, tetrahedral water, coexisting in aqueous solutions with varying populations(*36–43*). We note the selective expulsion of wrap to bulk water would be thermodynamically favorable due to overall entropy increase. If light activation is accompanied by selective expulsion of wrap water, this would thermodynamically favor and stabilize the lit state, and the apparent slowing of surface water dynamics can be due to an increased population of the slower diffusing, bound water and/or the eviction of the faster diffusing, wrap water into bulk water. The N414Q variant shows even greater degree of dehydration than WT according to ODNP data, suggesting that the N414Q mutation further stabilizes the lit state, in agreement with prior reports(*23*, *28*, *49*). We next seek direct experimental evidence using ^17^O NMR chemical shift analysis to support this claim.

### ^17^O NMR chemical shift shows distinct hydration water populations

The ^17^O NMR chemical shift of freely tumbling water is exquisitely sensitive to the electron density on the oxygen atom. It can detect subtle changes such as 0.01 Å differences in O-H covalent or hydrogen bond lengths and whether three or four hydrogens are bound to the central oxygen. In solution state under free tumbling conditions, the ^17^O NMR shift of water should not be altered significantly by the second-order quadrupolar effect. Hence it offers a direct experimental signature for different water populations(*63–69*). Persistent dynamic water structures in solution state has been recently demonstrated for lipid membrane surfaces by Zhang et al.(*70*) Furthermore, a series of studies of water/glycerol and water/DMSO mixtures identified bound and wrap water, categorized by differences in their tetrahedral content and reflected in the three body angles around 109°, to be the two dominant liquid water structural archetypes(*39*, *42*). These water structures have distinct hydrogen bond lengths, and hence different ^17^O chemical shifts, with wrap water with greater tetrahedral three-body hydrogen bond angles exhibiting a downfield shift relative to bound water(*71*). Bulk or hydration water contain both populations in varying proportions that are in dynamic exchange. The question is whether and how their makeup changes upon light activation of AsLOV2. To answer this question experimentally, we measured ^17^O NMR spectra of WT and slow-cycling N414Q solution prepared with 40% ^17^O enrichment water at 18.8 T and 294 K under MAS (10 kHz). The ^17^O *T*_1_ relaxation time of the solution containing WT AsLOV2 was 4.07 ms and N414Q was 5.15 ms. Both values were significantly shorter than the 7.05 ms value of pure water doped with the same ^17^O content, suggesting that hydration water populations with distinctly slowed dynamics are present in AsLOV2 solutions.

We used a pulse sequence established by Zhang et al (*70*) to suppress the ^17^O NMR signal from bulk water. Bulk ^17^O signal was minimized at the zero crossing (***τ***_zc_ = 2.70 ms) after an inversion pulse (Fig. 2A), so that the ^17^O NMR signal of protein-adjacent hydration water is enhanced relative to bulk water if their *T*_1_ is distinct from that of bulk water. With at 90°, the bulk-suppressed ^17^O NMR spectrum showed distinct signatures (Fig. 2C top, 2D top) up- and down-field of the bulk water peak. When varying the flip angle (θ) from 40° to 180° at fixed ***τ***_zc_ =2.7 ms and ***τ***_D_ = 0, we observed more effective suppression of bulk water signal at lower flip angles (θ < 100° for WT and θ < 130° for N414Q), manifested in more clearly resolved features of hydration water, consistent with observations by Zhang et al. As a control, we measured a 40% ^17^O enriched pure water sample under identical conditions (SI Fig. S4), where no distinct features were observed across the 0° to 180° range, confirming that the spectral signatures detected in the AsLOV2 samples arise from hydration water associated with the protein.

**Figure 2.**
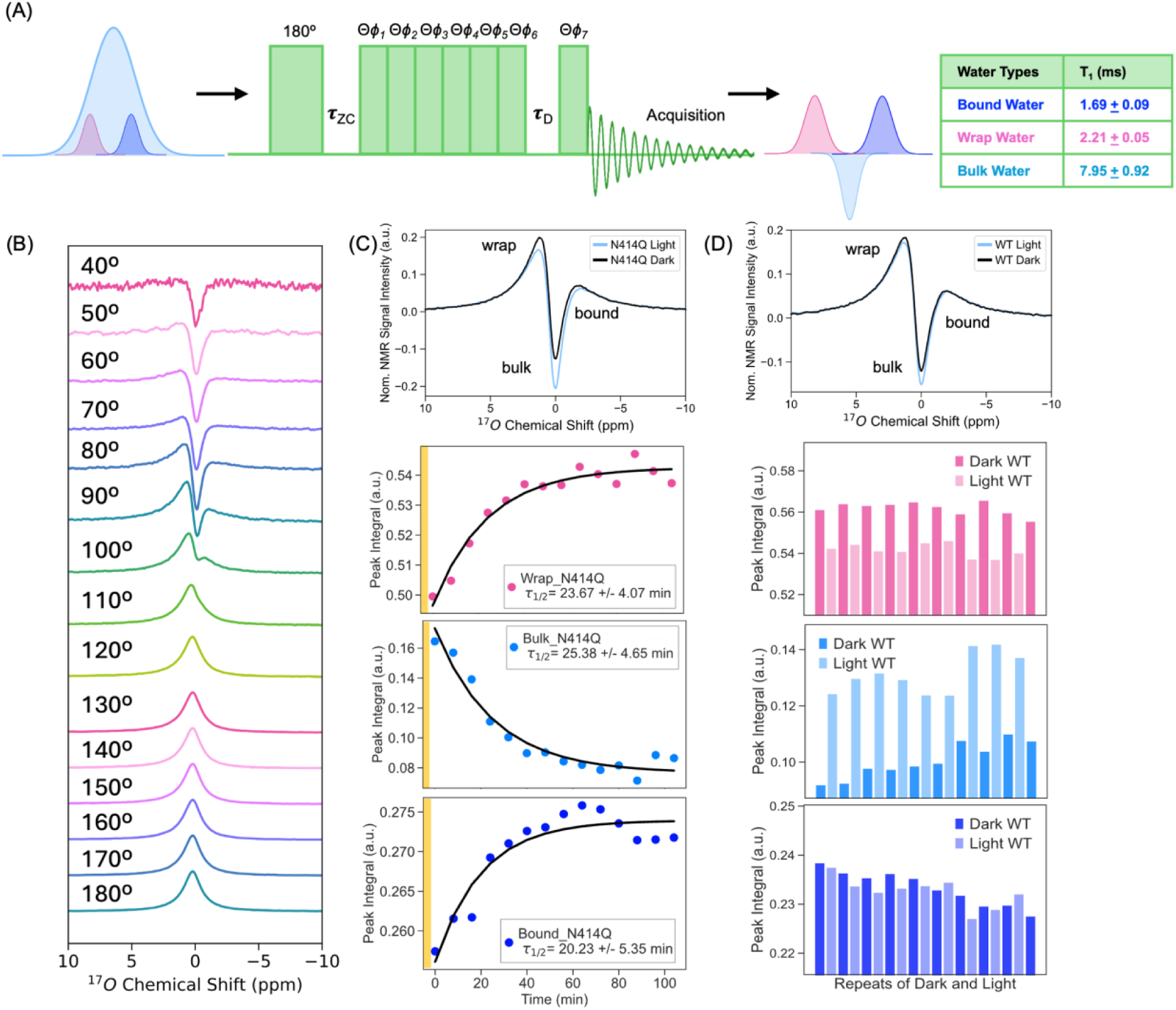
Light-Driven Redistribution of Distinct Water Populations in AsLOV2 Revealed by High-Field ^17^O MAS NMR. (A) Single-pulse 1D 17O MAS NMR spectrum of 40% 17O-enriched buffered AsLOV2 solution, acquired with bulk water suppression at 800 MHz and a spinning rate of 10 kHz ± 3 Hz at room temperature, reveals three spectrally distinct water populations. Bulk water exhibits the longest T_1_ (7.95 ± 0.92 ms), bound water shows the shortest T_1_ (1.69 ± 0.09 ms) and wrap water with intermediate T_1_ (2.21 ± 0.05 ms). (B) Experiments conducted with fixed τ_ZC_ =2.7 ms and τ_D_ = 0, while varying the flip angle θ, show that two water populations—assigned as “bound” and “wrap” water—are preferentially observed at low flip angles (<100°). (C) Due to its slower dark-state recovery, the N414Q mutant allows time-resolved tracking of hydration changes. Upon return to the dark state, the system exhibits gradual: recovery of wrap water, reduction in bulk water signal, and recovery of bound water, all occurring on a timescale of *τ*_1/2_ ~20 minutes, consistent with photocycle kinetics measured by time-resolved UV-Vis spectroscopy at 277 K. (D) In the wild-type protein, light activation leads to rapid and reversible: expulsion of wrap water, an increase in bulk water signal, and minimal change in bound water levels, as observed across repeated photocycles.

The greater spectral resolution at lower θ values was used to measure spectrally resolved *T*_1_ relaxation times (SI Fig. S4B) that reveal three discrete value ranges centered around: 2.21 ± 0.05 ms, 7.95 ± 0.92 ms and 1.69 ± 0.09 ms at ^17^O NMR shift ranges 8.0 to 0.5 ppm, 0.4 to −0.4 ppm and −0.5 to −8.0 ppm, respectively, relative to bulk water at 0 ppm. The spectral population exhibiting the longest *T*_1_ time, implying faster water dynamics also matches the *T*_1_ time of pure water of 7.05 ms, and was therefore assigned to bulk water corresponding to the 0.4 to −0.4 ppm range. The observation that distinctly shorter *T*_1_ values are found for the down- and up-field shifted spectral region suggests these contain protein-associated hydration water populations. Of these, we assign the down-field shifted signature as wrap water (2.21 ± 0.05 ms), consistent with DFT predictions for wrap water(*71*). The origin of the up-field shifted peak is less clear, but its shorter *T*_1_ (1.69 ± 0.09 ms) implies even slower dynamics than the down-field peak, and certainly contains protein-associated water, and potentially greater population of bound water. Since these populations of water are in dynamic exchange with each other in a dilute solution as our system, we anticipate the observed lineshape to be a convolution of spectral signatures of multiple populations that cannot be readily distinguished. Still, there are three clearly distinct spectral regions identified by different ^17^O NMR *T*_1_ values; hence we will focus on amplitude changes in each of these spectral regions upon light activation.

We next examined changes in the amplitudes of these three spectral regions upon activation of AsLOV2 with light. These measurements were conducted at a lower temperature of 277 K to slow the exchange rate between hydration water and bulk water, thereby enhancing the resolution and sensitivity for detecting light-induced changes in the hydration water composition. ^17^O NMR spectra with repeated activation shows that the light state induces a reversible decrease in wrap water and a concurrent increase in bulk water, with a minimal change to the up-field spectral population in WT (Fig. 2D). The fast recovery time of ***τ***_FMN_WT_ ~ 3.5 min (277 K) of WT did not allow for time-resolved measurements, given the acquisition time of 3 min for a ^17^O NMR spectrum. The slower dark state recovery time of N414Q of ***τ***_FMN_N414Q_ ~ 23 min allowed us to readily capture the recovery of wrap water and loss of bulk water population upon light activation as a function of time, yielding a time constant of ~20 min for all three populations, which is on par with the recovery time according to UV-Vis absorption at 277 K (Fig. 2C, S5). Time resolved measurements reveal that the up-field spectral population also changes, but to a much smaller extent than changes in the wrap and bulk water populations. Hence, changes in the ^17^O NMR chemical shift in solution state captures most prominently the light-induced expulsion of wrap water into bulk, with reversible recovery upon return to the dark state. Combined with ODNP results showing that protein-adjacent hydration water dynamics is slowed down by light activation, we conclude that the light activated state is hydrated to a greater extent with slow diffusing bound water upon eviction of the more dynamic, wrap water, in an entropy-driven process.

Our results also reveal that the change captured by ^17^O NMR is a collective eviction of wrap water upon light activation of AsLOV2, given that ^17^O NMR of water reports on changes of the entire population of water without site specificity. This gave rise to a compelling question: could the collective movement of hydration water induce the mechanical actuation of AsLOV2?

### Pressure to replace light as an inducer of AsLOV2 activation

If collective water movement induces conformational changes that accompany AsLOV2 activation, then any perturbation that induces a concerted eviction of structured hydration water could serve as a trigger. Naturally, this suggests pressure can replace light as an activator. We tested this hypothesis using a combination of computational and experimental methods. Computationally, we performed MD simulations of AsLOV2 at 3 kbar and compare to 1 bar to determine whether elevated pressure induces conformational changes consistent with light activation of AsLOV2 and eviction of wrap water. Unlike pressure activation, light activation involves changes in the electronic structure of FMN and surrounding molecules, and hence is outside the reach of classical MD. Experimentally, we performed DEER to measure distance change across spin labeled sites T406C-E537C upon light or pressure activation followed by vitrification, and 3D HNCO NMR in solution state to evaluate changes in the amide ^1^H-^15^N and carbonyl ^13^C chemical shifts of AsLOV2 residues under high pressure and compared these to changes upon light activation.

Proteins in solution state exist as an ensemble of conformational substates, each with distinct partial molar volumes. High pressure perturbs these equilibria by stabilizing conformations with reduced volume, often shifting populations toward folding intermediates or excited states that are otherwise sparsely populated under ambient conditions. While this concept of populating sparsely populated, excited states by high-pressure NMR(*72–76*) or EPR(*77–81*) has been established, the factors that govern intrinsic protein volume change and compressibility are only partially understood. We hypothesize that different populations of wrap and bound water constituting the hydration shell plays a critical role in pressure-induced conformational transitions: the lower density wrap water has greater compressibility and is preferentially displaced upon pressure increase, leaving behind the higher density hydration layer in which water remains more strongly bound to the protein surface. This transition may stabilize the protein’s excited state by increasing the total entropic contributions, minimizing the overall protein volume, and promoting solvation of newly exposed residues.

### Molecular dynamics simulations show expulsion of wrap water in pressure-activated AsLOV2

To test this hypothesis, we performed MD simulations of AsLOV2 at room temperature (298 K) under both ambient (1 bar) and elevated (3 kbar) pressure by computing transfer free energies(*82*) in implicit solvent for 200 ns following the protocol outlined by Arsiccio and Shea(*82*). Our results reveal a pressure-driven conformational transition consistent with light-induced activation, in which partial unfolding of AsLOV2 is initiated by the unraveling of the A’α helix, followed by extension and unfolding of the Jα helix, and the detachment of the Jα helix from the Iβ surface. The two helices move opposite to one another, with the Jα helix becoming more extended over longer equilibration times at high pressure (Fig. 3A, SI Fig. S14).

**Figure 3.**
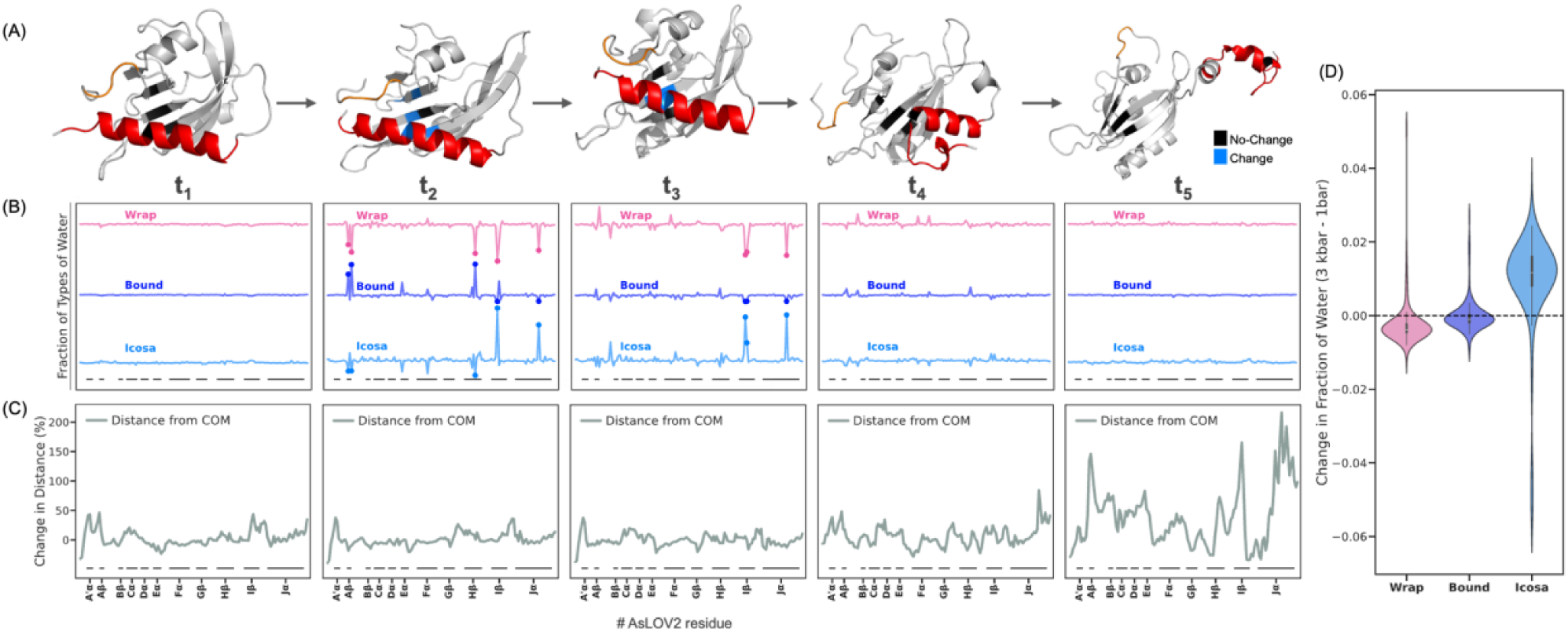
High-pressure molecular dynamics simulations reveal site-specific redistribution of water associated with pressure-induced unfolding in AsLOV2. (A) High-pressure molecular dynamics simulations reveal pressure-driven partial unfolding of AsLOV2, initiated by unraveling of the A’α helix (orange) and extension of the Jα helix (red), mirroring the structural changes observed upon photoactivation. (B) Snapshots from the 3 kbar unfolding trajectory reveal that wrap water remains stably associated with compact regions of the protein, such as at t_1_ (5% of total simulation time). By t_2_ (12% of total simulation time), a notable decrease in tetrahedral wrap water emerges around the Aβ, Hβ, Iβ, and Jα regions, accompanied by increased bound water in Aβ and Hβ and increased icosahedral water in Iβ and Jα. At t_3_ (27% of total simulation time) tetrahedral wrap water further decreases in Iβ and Jα, with a corresponding rise in icosahedral water. Jα begins to unfold at t_4_ (53% of total simulation time) and fully undocks by t_5_ (100% of total simulation time), where the wrap water redistributes to resemble the 1 bar state, though with slightly reduced overall occupancy, as shown in (D). The vertical positioning of the three plots is arbitrary. The wrap water values plotted here are scaled by 2.5X for clarity. Pink dots mark residues with >50% reduction in wrap water relative to the mean across all residues, while dark-blue and light-blue dots indicate bound and icosahedral water, respectively, which increase at residues losing wrap water at t_2_ ns and t_3_ ns. (C) Time-resolved residue-wise structural deviations relative to the 1 bar state reveal that pronounced changes begin to occur around t_5_ ns, particularly in the Iβ and Jα regions, suggesting that hydration loss precedes major conformational transitions. (D) Violin plot showing the overall pressure effect on local water populations: tetrahedral wrap water decreases, icosahedral water increases, while bound water remains nearly constant due to residue-specific behavior when pressurized from 1 bar to 3 kbar.

To determine whether pressure-induced unfolding involves the expulsion of tetrahedral like wrap water, we analyzed the water structure by computing the three-body-angle (3BA) formed by a central oxygen and two neighboring oxygens from hydrogen-bonded water molecules(*83–85*) using explicit-solvent simulations with the CHARMM36m force field and TIP3P-CHARMM water model(*86*). 196 representative unfolded conformations were obtained via RMSD clustering of the last half of 200 ns long unfolding trajectories following the method of Daura et al.(*87*) implemented in GROMACS(*88*), and change-point detection of Solvent Accessible Surface Area (SASAR) implemented using the Python ruptures package(*89*). Each unfolded conformation was then inserted into a new simulation box alongside explicit water and simulated for 500 ns. We examined the distribution of 3BA of water within the protein hydration shell and quantified the fraction of low-density tetrahedral water (3BA near 109.5°) at both 1 bar and 3 kbar partially unfolded states for the 200-500 ns simulation window. We define this tetrahedral water population as wrap water. We also calculated the residence times of hydration water (picoseconds) as a function of sequence position for AsLOV2 spanning residues 403–546 over the last 10 ns of production simulations at ambient and high pressure. Residence times were computed in first-passage mode, where re-entries are excluded from average lifetimes. Our analysis shows that closer to planar hydrogen-bond geometry (150°–170°) is associated with the longest residence times, suggesting a distinctive geometric motif for protein-bound water. We therefore classify this population as bound water (SI Fig. S11). Given that bulk water consists mostly of tetrahedral (100°–120°) and icosahedral water (50°–70°), and that tetrahedral water populations in the context is defined as wrap water, we probe the icosahedral water population as a representative of the remaining water species. This approach provides a more complete understanding of water structural transitions and enables us to monitor bulk-like water. As the protein unfolds, we found a consistent net decrease in the population of water with tetrahedral 3BA (i.e. wrap water), negligible net change in the population of bound water and a net increase in bulk-like icosahedral water populations (Fig. 3D, SI Fig. S12). For sites where significant change is observed, we see a decrease in tetrahedral wrap water for most sites, including the A’α and Jα helices, accompanied by concurrent increase in icosahedral water populations and/or increase in bound water populations (SI Fig. S12). This observation suggests that pressure-induced unfolding of AsLOV2 is closely associated with the eviction of tetrahedral wrap water consistent with experimental observation. This observation reveals a mechanism that is rarely considered: the compressibility of a protein depends on the structural property of protein-associated water. To date, protein compressibility has been conceptualized in terms of protein-driven determinants such as packing defects and cavities, while our findings demonstrate that modulating the population of waters of different density is a major factor(*74*, *90–92*). In fact, cavity- or solvation-based models for compressibility may be strongly correlated. Future studies will have to investigate whether wrap water eviction is a general mechanism in high pressure-mediated excited state populations.

### DEER studies show that light and pressure are additive factors for AsLOV2 activation

Next, we experimentally compare effects of both light and pressure (3 kbar)(*77–81*) on the conformational changes in AsLOV2 using DEER of doubly spin labeled WT and N414Q AsLOV2 at sites T406C and E537C. DEER measurements in the dark state revealed an inter-site distance distribution, *P*(*r*), centered around 2.2 nm in both variants, while light activation results in an extension in WT and N414Q by 1-3 nm (Fig. 1A, C). Application of 3 kbar pressure alone (see SI for method) did not result in a measurable shift from this equilibrium distance in the WT, while ~3% of the N414Q adopted the extended population with distance centered around ~4 nm (Fig. 4B, F). In contrast, light activation alone resulted in ~10% of the WT and ~9% of the N414Q population adopting an extended conformation (Fig. 1A, C). When both stimuli, light and pressure, were applied simultaneously, the effects were additive, with 21% of the WT and 26% of the N414Q population adopting the extended conformation, surpassing the effect of either stimulus alone (Fig. 4B-G). Given that our experimental setup was limited to 3 kbar, we increased the internal osmotic pressure using a molecular crowder (PEG 20 kDa) to examine the effects of higher pressure. PEG alone induced a modest shift, with ~3% of the WT and ~4% of the N414Q population adopting an extended conformation. When PEG was combined with a hydrostatic pressure of 3 kbar, the population in the extended state of WT increased to 4% and of N414Q to 7%, suggesting a small, but measurable additive, effect. The combination of light, pressure, and PEG yielded the largest conformational shift, with 26% of the WT and 79 % of the N414Q adopting the extended conformation (Fig. 4C, E, F, G).

**Figure 4.**
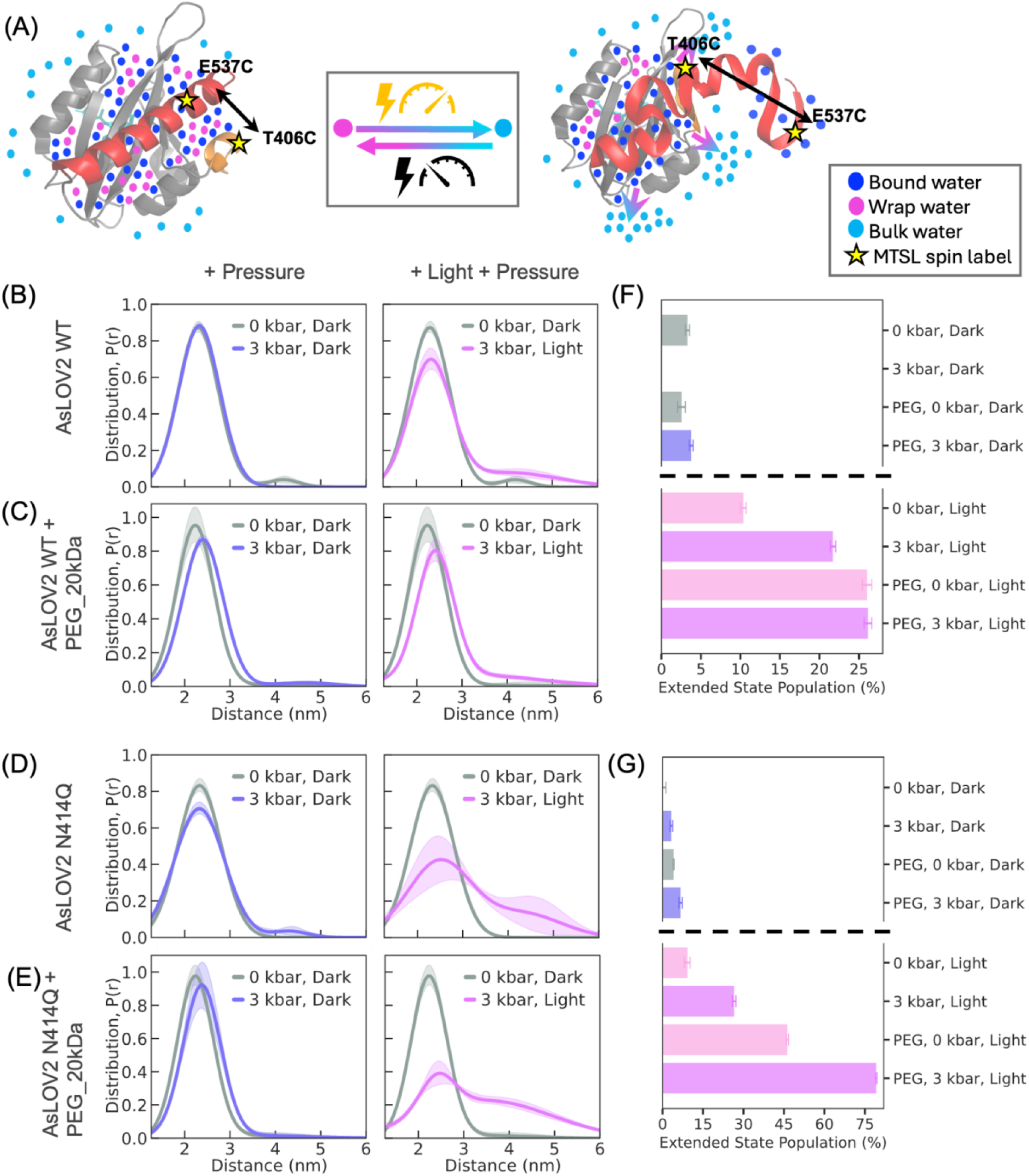
DEER measurements show that pressure and light co-operatively enhance the population of extended state conformations in AsLOV2, with effects amplified by molecular crowding (see SI for details of the analysis of the DEER data). (A) Schematic of MTSL spin-labeled AsLOV2 at positions T406C and E537C, illustrating conformational extension and concurrent release of wrap water upon stimulation by light and/or pressure. (B) Left: DEER-derived distance distribution *P*(*r*) shows that application of 3 kbar pressure alone did not result in a detectable structural extension from the dark-state equilibrium conformation in the WT. Right: Combined light and pressure further increase the extended population to ~21%, highlighting an additive effect. (C) In the presence of PEG_20kDa as a molecular crowder, pressure alone increases the WT extended population to ~4% (left), and light plus pressure drives it further to ~26% (right), suggesting enhanced conformational responsiveness due to internal osmotic pressure. (D) In the N414Q mutant (no crowder), pressure alone results in ~3% of the population in the extended conformation (left), while light plus pressure increases this to ~26% (right), indicating greater stabilization of the lit state. (E) In the presence of PEG_20kDa, the N414Q mutant shows even stronger pressure sensitivity: ~7% in the extended state under pressure alone (left), and ~79% with combined light and pressure (right). (F–G) Summary of population shifts under different conditions, indicating that both external (hydrostatic) and internal (osmotic) pressure cooperatively enhance conformational extension in AsLOV2, with the N414Q mutant exhibiting a greater propensity for adopting the excited-state conformation.

Notably, the effect of pressure on the slow-cycling N414Q variant is much greater, suggesting that the N414Q variant stabilizes the excited state more effectively than the WT. In light of the ODNP results showing that N414Q undergoes greater extent of dehydration, the observation that pressure activation has a greater effect on stabilizing the excited state of N414Q compared to WT suggests that N414Q is more readily compressible and that dehydration stabilizes the excited state of N414Q. These results provide experimental evidence that both pressure and light induce conformational extension and dehydration of AsLOV2 might be a common mechanism. The findings that a molecular crowder such as PEG shows additive effects to light and pressure supports our hypothesis that hydraulic action is at the core of AsLOV2 activation. This study demonstrates that nature might be leveraging both external and internal pressure to regulate functional conformational transitions in LOV domains, rendering them mechanosensitive.

### HNCO NMR shows pressure activation induces similar conformational changes as blue light

The DEER results showing pressure-induced conformational extension between the A’α and Jα helices motivated an inquiry into structural changes at residue-level resolution across the entire protein to offer an experimental complement to the high-pressure MD results. To accomplish this, we employed high-pressure three-dimensional (3D) HNCO (¹H/¹⁵N/¹³C) NMR spectroscopy on AsLOV2, across a pressure range up to 2.5 kbar using a high-pressure ceramic tube (Daedalus Innovations), to measure pressure-induced chemical shift perturbations in ¹H, ¹⁵N, and ¹³C nuclei of the protein backbone. Representative two-dimensional (2D) ¹⁵N/¹H projections from the 3D NMR dataset illustrate individual resonances undergoing either linear or nonlinear chemical shift changes with increasing pressure (Figure 5A). To systematically quantify these shifts across all three nuclei, we performed least-squares fitting of pressure-dependent chemical shifts for each residue to a quadratic function of pressure:

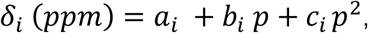

where *δ_i_* is the chemical shift (ppm) for the *i*-th residue at pressure *p* (bars), *a_i_* is the chemical shift at 20 bar, and *b_i_* and *c_i_* are the linear and nonlinear coefficients, respectively. This empirical model is analogous to the thermodynamic expansion of Gibbs free energy as a function of pressure, allowing for a biophysical interpretation of the coefficients: *b_i_* corresponds to changes in the partial molar volume, while *c_i_* captures changes in compressibility of site *i*.

**Figure 5:**
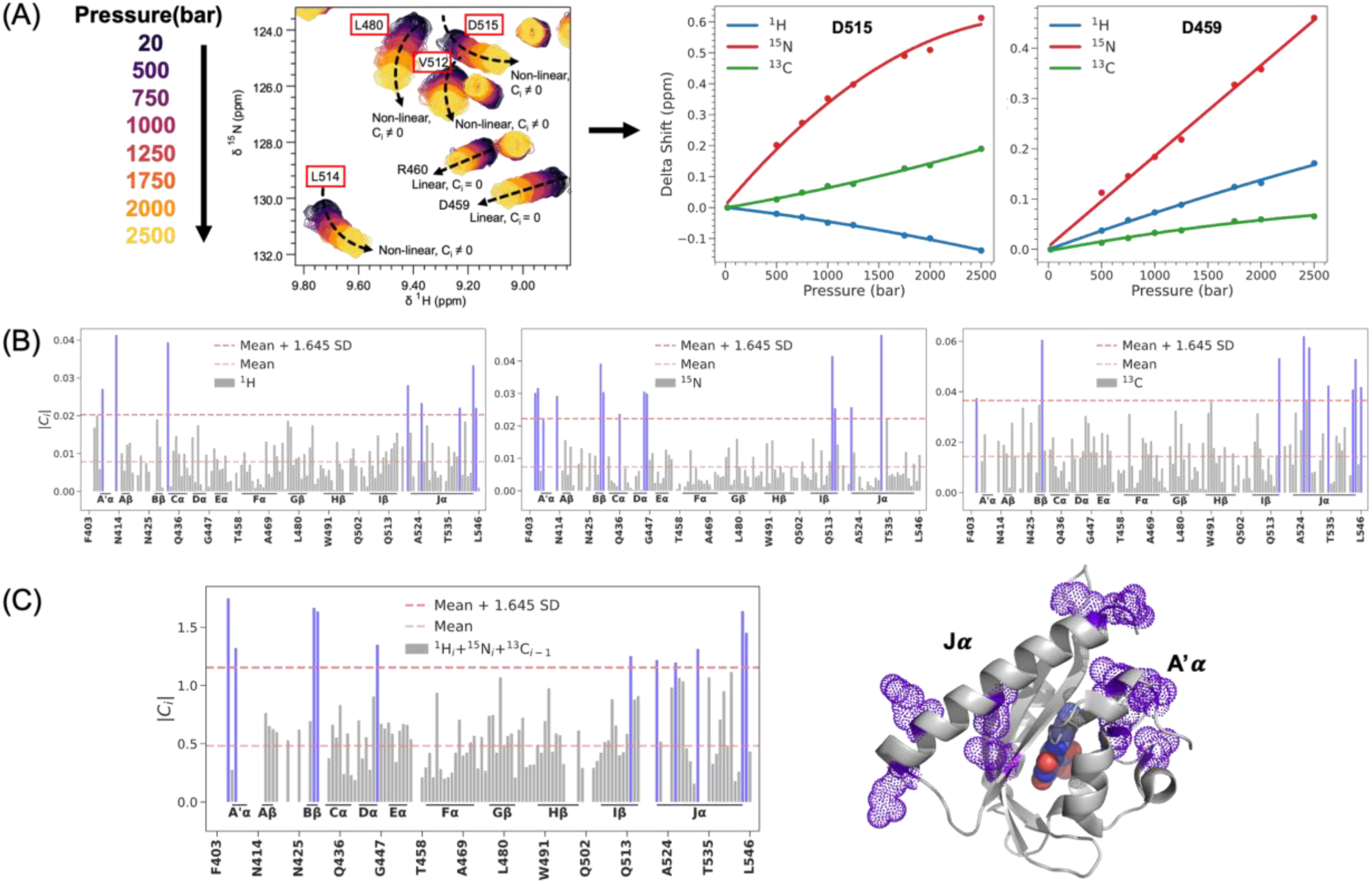
Residue-resolved analysis of pressure-induced conformational changes in AsLOV2 using high-pressure 3D HNCO NMR. (A) Representative two-dimensional ¹⁵N/¹H projections from high-pressure HNCO spectra collected across a pressure range up to 2.5 kbar. Peak trajectories illustrate pressure-dependent chemical shift changes, including both linear and nonlinear behaviors across different residues. (B) Quadratic fitting of chemical shift changes for three backbone nuclei—¹H, ¹⁵N, and ¹³C—was used to extract nonlinear coefficients (*c_i_*) for each residue. Bar plots show the distribution of absolute nonlinear coefficients |*c_i_*| across the protein for each nucleus, with horizontal lines indicating the statistical cutoffs (mean, and mean + 1.645 SD). (C) A composite nonlinearity score was calculated by normalizing and summing the nonlinear coefficients across all three nuclei (¹H*_i_* + ¹⁵N*_i_* + ¹³C*_i-1_*), yielding an integrated metric for pressure sensitivity per peptide group. Structural mapping of the top 10% of residues with the largest deviations from the mean onto the AsLOV2 crystal structure (PDB: 2V1A (*62*)) reveals that the most pressure-sensitive regions are localized to the A′α and Jα helices and their adjacent segments. These helices are also known to be functionally critical in the light activation mechanism of AsLOV2, suggesting a shared structural pathway for both light- and pressure-induced activation via hydration-mediated “hydraulic” coupling.

Residues exhibiting linear pressure shifts (*c_i_* = 0) are interpreted as undergoing population redistributions among conformational substates within the native ensemble, usually due to isotropic compression of the protein by the applied pressure. In contrast, residues with significant nonlinear components (*c_i_* ≠ 0) report on a population shifts toward pressure-favored conformations with altered compressibility that correspond to a new state not populated at equilibrium under ambient pressure(*72–76*). To identify residues undergoing the most pronounced conformational changes, we calculated the absolute value of the nonlinear coefficient (*c_i_*) for each residue and applied statistical thresholds based on mean and standard deviation (SD) of 1.645. This threshold captures the top 10% of residues with the largest deviations from the mean, corresponding to major sites experiencing significant pressure-induced changes. (Figure 5B).

The chemical shift behavior of different nuclei offers distinct insights: ^1^H shifts correlate strongly with changes in the H^…^O hydrogen bond distance in the amid bond motifs; whereas the ^15^N shifts are sensitive to variations in backbone torsion angles (*72–76*). In contrast, ^13^CO chemical shifts, which have been less commonly studied experimentally, report on both parameters but offer a more global structural readout of changes in the secondary structure. To integrate the complementary information provided by each nucleus and obtain a comprehensive residue-level measure of nonlinear response, we normalized the nonlinear coefficients across the three nuclei and calculated a combined nonlinearity score for each peptide group by summing the normalized values. This integrated metric provided a holistic, residue-level assessment of pressure-sensitive conformational changes throughout the protein backbone (Figure 5C).

The top 10% of residues experiencing pressure-sensitive conformational changes lie primarily within A′α (T406, L408), Bβ (F429, A430), Dα (L446), Iβ, (L514), and Jα (R521, R526, I532, K544, E545) segments. These findings show that the N- and C-terminal regions of AsLOV2 undergo substantial pressure-induced conformational changes. Notably, these regions correspond to critical structural elements involved in the blue-light activation mechanism(*22*). This concordance implies that high pressure does not indiscriminately destabilize the protein structure but activates AsLOV2 by populating excited structural states as light does. We propose a common hydration-mediated “hydraulic” coupling mechanism underlying both pressure- and light-mediated activation. This conclusion is further reinforced by TiGGER and DEER results, which independently revealed related conformational changes that drive the A’α and Jα helices apart by several nanometers, implying that the Jα helix undocks from the Iβ surface, induced by additive effects of pressure and light activation.

### Does hydraulic action follow or precede protein activation?

The next critical question is whether the concerted water action precedes or follows conformational changes. The observation that pressure can induce comparable and additive activation as light, and that low-density wrap water populations are selectively evicted upon light activation (according to ODNP and ^17^O NMR) and pressure activation (according to MD), suggests that the common driver is hydraulic action that drives protein activation. However, with the current technology available, direct access to the sequence of events of protein and water movement of AsLOV2 under physiological condition is only possible computationally, informing our quest whether pressure-induced eviction of wrap water precedes the structural transformation of AsLOV2.

To investigate the temporal series of changes in conformational changes and water, we analyzed multiple time points along the high-pressure (3 kbar) MD trajectory. The A’α helix undergoes complete unfolding within the first 2.5% of total simulation time (SI Fig.S13), yet the overall distribution of wrap water remains largely unchanged across residues up to t_1_ at 5% of total simulation time. At around t_2_ at 12% of total simulation time we observe a sudden and significant decrease in tetrahedral wrap water surrounding residues in the Aβ (V416, T418), Hβ (L496), Iβ (I510), and Jα (A536) regions, while the Jα helix and Hβ and Iβ fold remain fully intact. This loss in wrap water coincides with an increase in bound water in Aβ and Hβ and an increase in bulk-like icosahedral water in Iβ and Jα. By t_3_ at 27% of total simulation time, further eviction of wrap water is observed, but more pronounced around the Iβ (I510, G 511) and Jα (A536) domains, while the Jα helix is still structurally fully intact. After t_4_ at 53% of total simulation time, no more significant changes in local hydration are detected, while first signs of Jα helix unfolding emerge at site A536. Finally, by t_5_ at 100% of total simulation time, an overall reduced population of wrap water redistributed across all sites is observed, while the Jα helix has fully undocked and moved away from the A’α helix, (Fig. 3B). This analysis shows that eviction of wrap water occurs between t_1_ and t_3_. To assess the magnitude of global structural change at each time point, we computed residue-wise displacement relative to the center of gravity of the initial 1 bar structure. Minimal fluctuations were observed across the protein up to t_4_, consistent with the stability of key secondary structure elements observed. Only around t_5_, pronounced global conformational changes emerge, particularly in the A’α, Iβ and Jα regions, confirming that changes in hydration precede the major conformational transitions in AsLOV2 (Fig. 3C).

## Discussion and Conclusion

This study debuts a model for protein activation for AsLOV2 in which light or pressure-induced unfolding follows the concerted loss of low-entropy, tetrahedral, hydration water, particularly near the critical Jα helix. What began as an exploration to capture light-driven conformational extension of AsLOV2 evolved into the recognition of a novel mode of protein activation by a concerted and reversible hydraulic action of water moving in and out of the protein. Using a combination of time-resolved EPR and DEER spectroscopy, we observed a long-range structural extension upon blue-light illumination of the tip of the Jα helix relative to A’α, a hallmark of functional LOV activation. ODNP experiments revealed a surprising slowdown in surface water diffusivity upon light activation, even at highly water-accessible protein surface sites, hinting at a light-triggered restructuring of the hydration shell of AsLOV2. High field ^17^O NMR allowed us to spectroscopically resolve the distinct populations of bulk, wrap, and bound water, and crucially, track their collective and reversible redistribution during photoactivation, and validating our hypothesis that wrap water is selectively evicted, transforming into bulk water, and recalled to hydrate AsLOV2 in the dark state. Molecular dynamics simulations under elevated pressure not only recapitulated the light-induced conformational transition but revealed pressure as an alternative activator for AsLOV2, acting through the same mode of expulsion of low-density, tetrahedral wrap water. These predictions were experimentally confirmed by high-pressure DEER, which showed light- and pressure-induced extensions, further amplified by increasing internal pressure by molecular crowding. High-pressure 3D HNCO NMR uncovered residue-specific nonlinear responses concentrated in helices known to mediate light signaling, reinforcing the connection between hydration modulation and structural activation. Lastly, MD simulations revealed that concerted water eviction precedes protein conformational extension, offering further support for the thesis of hydraulic activation of AsLOV2.

AsLOV2 as a hydraulic actuator is a novel conceptual framework, distinct from earlier models that viewed water as a molecular constituent serving as a proton donor or acceptor. The concept that water serves as a hydraulic fluid to exert mechanical work in a thermodynamic process changes the way we understand, modify and model signal transduction in AsLOV2. MD simulations can be used to discover mutations that stabilize high-pressure induced excited state conformations and model the mechanical force produced by these conformational changes. We offer a testable hypothesis: hydraulic activation is a generalized mechanism of mechanosensitive LOV domains that produces mechanical work upon light activation. This concept provides a new lens for understanding protein responsiveness to both optical and mechanical input. The discovery that pressure, like light, can drive this transition has implications for designing externally controllable protein-based systems. This is supported by demonstrations of protein-based switches existing in off states at equilibria that are turned on by specific stimuli, e.g. light with AsLOV2 (*93*), but also by point mutations (*94*) and general thermodynamic variables such as temperature (*95*). Our work extends this to an often-underused parameter – pressure – and offers a new mechanistic framework of hydraulic action underpinning protein activation. This study advances the blueprint for rationally engineering and testing optogenetic tools and to photoacoustic actuators whose activity can be finely tuned for applications in synthetic biology, smart materials, and targeted therapeutic modulation.

While our study provides converging evidence for a hydration-mediated activation mechanism in AsLOV2, several opportunities remain to sharpen and expand this framework. Time-resolved methods such as pressure-jump NMR, EPR and diffraction-based methods will help resolve the temporal sequence of hydration change and conformational transition events. Additionally, whether pressure-induced eviction of wrap water is independent of chromophore chemistry remains to be explored. We speculate that pressure populates the same spin states for FMN as light, but this requires unprecedented experimental capabilities of time-resolved EPR under high pressure that should be subject of future studies. The combination of electronic structure calculation and atomically detailed MD simulations may offer more direct insight into light-activated changes in protein-associated water and offer more quantitative insight into the hydrogen-bonding geometries and electronic environments that distinguish wrap, bound, and bulk water populations, and how the transient triplet state population of FMN induces hydraulic expulsion of tetrahedral water populations. Extending the here presented studies to other LOV proteins will be critical to determine whether hydraulic activation is a broadly conserved strategy or a finely tuned adaptation, offering deeper insight into how proteins use water not just as solvent, but as an active medium for transducing environmental signals into mechanical motion.

## Acknowledgements

We acknowledge support from NSF grants MCB-2025860 and CHE-2411584 for biophysical studies of AsLOV2 using UV-Vis spectroscopy and TiGGER. Support for ODNP experiments comes from the NIH MIRA grant R35GM136411 and the Deutsche Forschungsgemeinschaft (DFG, German Research Foundation) as a part of Germany’s Excellence Strategy EXC2033 390677874 RESOLV. Hydration studies using ^17^O MAS NMR made use of the MRL Shared Experimental Facilities supported by the MRSEC Program of the National Science Foundation under Award No. DMR 2308708. We thank Ms. Kierson Rickmon for assistance with AsLOV2 expression and purification. We are grateful to Dr. Jerry Hu and Ms. Jaya Nolt for their expert assistance and for overseeing operations at the MRL Shared Experimental Facilities. High-pressure solution-state NMR experiments were conducted at the CUNY ASRC Biomolecular NMR Facility, with support from NIH grants R01 GM106239 (to K.H.G.), R35 GM156296 (to K.H.G.), and R01 GM123012 (to B.A.J.). We thank Dr. Denize C. Favaro for excellent technical assistance and for her support of the CUNY ASRC Biomolecular NMR Facility. DEER spectroscopy studies of conformational change in AsLOV2 were conducted at the University of St. Andrews, supported by EPSRC (studentship to HR EP/R513337/1), BBSRC (BB/T017740/1) and Wellcome Trust (099149/Z/12/Z) for the Q-band EPR spectrometer and latterly also the high-pressure equipment. We thank Drs Hassane El Mkami, Robert Hunter, David Norman, and Mr Callum Smith for technical assistance. J.-E.S. acknowledges support from the NSF (MCB-1716956). Simulations were supported by the NSF grant ACI-1548562 (project TG MCA05S027 using the Purdue Anvil Cluster), from the Advanced Cyberinfrastructure Coordination Ecosystem: Services & Support (ACCESS) program, which is supported by U.S. National Science Foundation grants #2138259, #2138286, #2138307, #2137603, and #2138296, and the computational facilities purchased with funds from the National Science Foundation (CNS-1725797) and administered by the Center for Scientific Computing (CSC). The CSC is supported by the California NanoSystems Institute and the Materials Research Science and Engineering Center (MRSEC; NSF DMR 2308708) at UC Santa Barbara.

