## Supplemental Information for "Hydraulic Activation of the AsLOV2 Photoreceptor"

### 1. Protein expression

A DNA fragment encoding the LOV2 domain of Avena sativa phototropin 2 (residues 403–546) was purchased from GENE UNIVERSAL INC., cloned into the pET-26b(+) vector via NdeI (5′) and XhoI (3′)

restriction sites, and included a C-terminal 6×His affinity tag. The plasmid was transformed into E. coli BL21 (DE3) cells and plated on LB agar containing kanamycin (0.05 mg/mL).

Primary culture: A single colony was used to inoculate 10 mL LB medium supplemented with kanamycin (0.05 mg/mL), which was grown at 37 °C with shaking at 220 rpm for 18 hours.

Secondary culture: The overnight culture (primary culture) was then used to inoculate 1 L of LB medium containing kanamycin and grown under the same conditions until reaching an optical density (OD600) of 0.7–0.8. Protein expression was induced with 1 mM IPTG, and cultures were incubated for ~16 hours at 18 °C in the dark with continuous shaking.

> For U- $^{15}$N-$^{13}$C labeled samples (for high pressure NMR): The overnight culture (primary culture) was used to inoculate 1 L of LB medium containing kanamycin and grown under the same conditions until reaching an optical density (OD600) of 0.8–1.0. Cells were harvested by centrifugation and resuspended in 250 mL of M9 minimal medium supplemented with $^{15}NH_4Cl$ (1 mg/mL) as the sole nitrogen source and [U-$^{13}C_6$] glucose (3 mg/mL) as the sole carbon source. The culture was incubated for an additional 30 minutes at 37 °C with shaking (220 rpm) to allow metabolic adaptation. Protein expression was then induced with 1 mM IPTG, and cells were incubated for ~16 hours at 18 °C in the dark with continuous shaking.

Cells were harvested by centrifugation at 5,000 × g for 15 min at 4 °C and resuspended in 30 mL of lysis buffer (20 mM Tris-HCl, 500 mM NaCl, pH 7.8) containing 5 mM β-mercaptoethanol and lysozyme (1 mg/mL). After a 30-minute incubation at 4 °C, cells were lysed by sonication (60 cycles of 10-second pulses with 10-second intervals) and clarified by centrifugation at 11,000 rpm for 30 min at 4 °C to remove cellular debris.(*1*)

### 2. Protein purification

To minimize the population of apo-protein, exogenous flavin mononucleotide (FMN) was added to the lysate at a 150-fold molar excess relative to the expected protein concentration. The mixture was incubated at room temperature in the dark under gentle rotation (15 rpm) for 30 minutes. For affinity purification, 5 mL of Ni-NTA resin was added, and the mixture was further incubated at 4 °C under rotation (15 rpm) in the dark for 2 hours. The suspension was then allowed to settle by gravity, and the supernatant was carefully removed. The resin-bound protein was washed three times with 40 mL wash buffer (20 mM Tris-HCl, 500 mM NaCl, 40 mM imidazole, 5 mM β-mercaptoethanol, pH 8.0). The protein was eluted using 10 mL of elution buffer (20 mM Tris-HCl, 500 mM NaCl, 500 mM imidazole, 5 mM β-mercaptoethanol, pH 8.0) by gravity flow through a chromatographic column. The eluate was subjected to size-exclusion chromatography using a HiLoad™ 16/600 Superdex™ 75 pg column (GE Healthcare, Chicago, IL) connected to an NGC™ Medium-Pressure Liquid Chromatography System (Bio-Rad, Hercules, CA). The column was equilibrated with SEC buffer (20 mM Tris-HCl, 150 mM NaCl, pH 7.5), and the protein was eluted under isocratic conditions. The pooled fractions were concentrated to a desired amount for further experiments.(*1*)

### 3. Protein spin-labeling

For site-directed spin labeling, purified AsLOV2 variants (L437C, V478C, L514C, N414Q-L514C, N414Q-T406C/E537C, E537C, and T406C/E537C) were first incubated with 10 mM dithiothreitol (DTT) at 4 °C overnight to fully reduce cysteine thiol groups. Following reduction, DTT was removed using a PD-10 desalting column (GE Healthcare, Chicago, IL), and the protein was immediately reacted with a 10-fold molar excess of spin label. Two spin labels were employed: the nitroxide-based MTSL (1-Oxyl-2,2,5,5-tetramethyl-Δ³-pyrroline-3-methyl methanesulfonothioate) for Overhauser Dynamic Nuclear Polarization

(ODNP) and double electron-electron resonance (DEER), and the $Gd^{3+}$-chelating probe Gd-sTPATCN for time-resolved Gd-Gd EPR (TiGGER) measurements. Labeling was carried out at 4 °C under gentle shaking overnight in the dark. Excess unreacted spin label was removed by size-exclusion chromatography using a HiLoad™ 16/600 Superdex™ 75 pg column (GE Healthcare, Chicago, IL) on an NGC™ Medium-Pressure Liquid Chromatography System (Bio-Rad, Hercules, CA), equilibrated with 20 mM Tris-HCl, 150 mM NaCl, pH 7.5. Final spin-labeled AsLOV2 samples were concentrated to ~1.5 mM for TiGGER and ~400 μM for ODNP experiments. For DEER experiments, spin-labeled proteins were further buffer-exchanged into deuterated buffer (20 mM Tris-HCl, 150 mM NaCl, pH 7.5 in $D_2O$) and concentrated to ~80 μM prior to measurement. Protein concentration was determined by absorbance at 447 nm using molar extinction coefficient of 13,500 $M^{-1}$ $cm^{-1}$.(*1*)

### 4. X-band Continuous Wave (CW) EPR Spectroscopy

CW EPR spectra were acquired at 295 K using a Bruker EMX X-band spectrometer equipped with a dielectric cavity featuring an optical window (model ER 4123D). Samples were photoexcited using a 450 nm blue laser (Laser Components USA, Inc., Bedford, NH) delivering 70 mW of output power, coupled into an optical fiber that transmitted approximately 15 mW to the sample space. Samples (10 μL) were loaded into quartz capillaries with an inner diameter of 0.66 mm (VitroCom). Spectra were collected first in the dark and subsequently under continuous 450 nm illumination. Measurements were conducted at a center magnetic field of 0.35 T and a microwave frequency of 9.8 GHz. Spectra were recorded with signal averaging over 10 scans, using a microwave power of 2 mW, a modulation amplitude of 0.5 G, a modulation frequency of 100 kHz, and a magnetic field sweep width of 150 G.

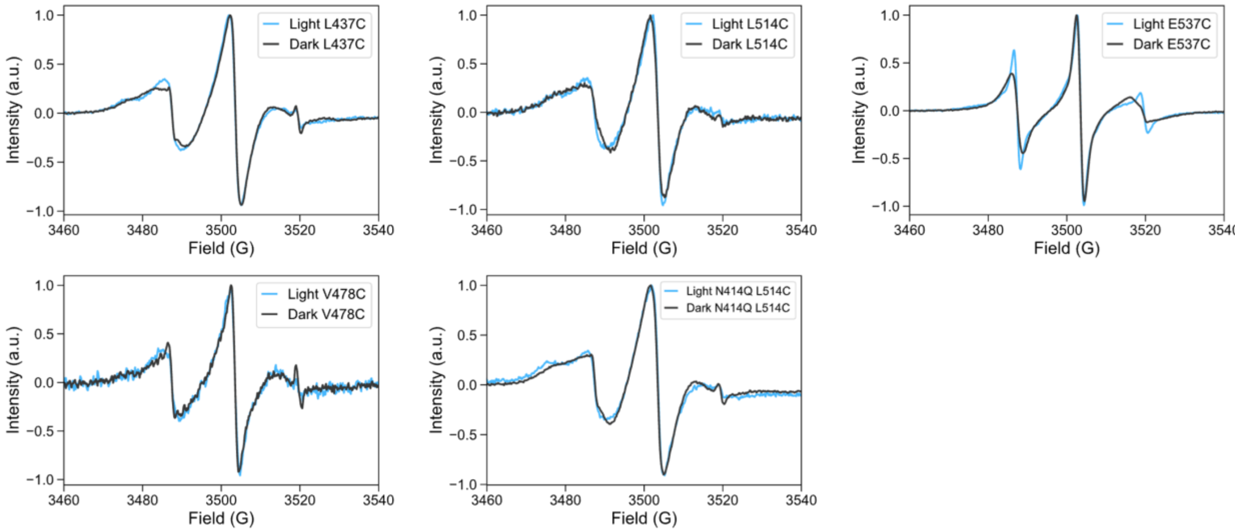

**Figure S1**. X-band continuous wave (CW) EPR spectra of AsLOV2 singly labeled with MTSL at L437C, V478C, L514C, L514C(N414Q), and E537C, measured at room temperature in the dark state (black) and with blue-light activation (blue). Most sites exhibit minimal changes in lineshape upon light activation. In contrast, site E537C displays clear motional narrowing in the lit state, consistent with increased side-chain dynamics upon Jα helix unfolding.

### 5. Overhauser dynamic nuclear polarization (ODNP)

#### 5.1. Experimental Setup

Overhauser dynamic nuclear polarization (ODNP)–enhanced $^1$H NMR experiments were performed using the same Bruker EMX X-band EPR instrument setup described above, except with a Bruker cavity with optical window (model ER 4119 HS) and a Bridge12 NMR detection coil (model: B12T ODP 9GHz B12TODP0019 R3) positioned within the cavity. The coil was tuned to 14.95 ± 0.1 MHz using a vector network analyzer and connected to the broadband channel of a Bruker Avance 300 NMR spectrometer. Magnetic field homogeneity was optimized by shimming the electromagnet using a set of Bridge12 shim coils (model: E-Shims B12TAES0002). Microwave irradiation was supplied by a Bridge12 microwave source (model: MPS 9GHz B12TMPS0006). Data acquisition was controlled using previously established methods implemented via a custom Python script (package: pyB12MPS), which initiated the NMR pulse sequence by sending a trigger from the microwave bridge to the NMR console. $^1$H signal enhancement curves were acquired by incrementally increasing the microwave power from 0 to 33 dBm over 19 steps. Spin-lattice relaxation times ($T_1$ and $T_{100}$) were measured using a standard inversion-recovery pulse sequence with optimized delay times and calibrated 90° excitation pulses.

#### 5.2. Data-Processing

The efficiency of coupling between the electron spin and nearby water protons is quantified by the coupling factor $\xi$, which encodes key information about local hydration dynamics. It is defined by the ratio of two fundamental relaxivity parameters: $k_\sigma / k_\rho$

The cross-relaxivity $k\sigma$ is sensitive to fluctuations of the electron–proton dipolar interaction at frequencies on the order of tens of GHz, and therefore selectively reports on fast, picosecond-scale dynamics. In contrast, $k_\rho$ includes contributions from both high-frequency (ps-scale) and low-frequency (ns-scale) fluctuations, the latter corresponding to the NMR Larmor frequency (~15 MHz). By subtracting the high-frequency contribution (i.e., the component reflected by $k_\sigma$) from $k_\rho$, we isolate the low-frequency relaxivity component $k_{low}$, which is exclusively sensitive to nanosecond-scale motions.

The degree of polarization transfer from electron spins to nuclear spins, expressed as the $^1$H NMR enhancement ($\varepsilon(p)$), was calculated using the following relation:

$$\varepsilon(p) = \frac{I(p)-I(0)}{I(0)}$$

where $I(p)$ represents the integrated intensity of the water $^1$H NMR peak acquired in the presence of microwave power $p$. The enhancement factor $\varepsilon(p)$ is a unitless quantity, defined relative to the NMR signal acquired in the absence of microwave irradiation, $I(0)$.

The value of the cross-relaxivity parameter $k\sigma$ were derived from the experimentally obtained values of $\varepsilon(p)$, in combination with longitudinal relaxation time ($T_1(p)$) measurements:

$$k_\sigma s(p) = (\frac{|\omega_H/\omega_e|}{C_{SL}}) e(p) T_1^{-1}(p)$$

Here, $T_1(p)$ denotes the longitudinal relaxation time of the $^1$H NMR signal measured at each microwave power $p$. The saturation factor $s(p)$ quantifies the net saturation of the electron spin transitions across all three hyperfine components of the nitroxide radical at a given microwave power. $C_{SL}$ is the concentration of the spin label, and $\omega_H$ and $\omega_e$ are the Larmor frequencies of the proton and electron, respectively.

The resulting $k_\sigma s(p)$ values were fit to the following saturation model and extrapolated to infinite microwave power:

$$k_\sigma s(p) = \frac{k_\sigma s_{max} p}{p_{1/2}+p},$$

where $p$ is the applied microwave power, $p_{1/2}$ is the power at which the electron spin transition reaches half-saturation, and $k_\sigma s_{max}$ is the maximum product of cross-relaxivity and saturation factor. It has been shown that for tethered nitroxide spin labels, $s_{max} \approx 1$, and thus $k_\sigma \approx k_\sigma s_{max}$ provides an excellent approximation for the cross-relaxivity.

To determine $k_\rho$, a control sample was prepared under identical conditions and at the same concentration as the spin-labeled sample, but without the spin label. The longitudinal relaxation time of this control sample, $T_{100}$, was measured using an inversion-recovery NMR experiment in the absence of microwave irradiation. This value was then compared to the $T_{10}$, relaxation time of the corresponding spin-labeled sample, also acquired without microwave irradiation. The relaxivity parameter $k_\rho$ was then calculated according to:

$$k_\rho = \frac{1}{C_{SL}}\left(\frac{1}{T_{10}} - \frac{1}{T_{100}}\right),$$

where $C_{SL}$ is the spin label concentration.

From the derived values of $k_\sigma$ and $k_\rho$, the self-relaxivity parameter $k_{low}$ was determined using the following equation:

$$k_{low} = \frac{5}{3}k_\rho - \frac{7}{3}k_\sigma$$

Finally, the dipolar coupling factor $\xi$, which is sensitive to local hydration dynamics, depends on the dipolar correlation time $\tau_c$ that characterizes the timescale of electron–proton dipolar interactions. This dependence is mediated through a spectral density function, $J(\omega;\tau_c)$, and can be quantitatively modeled using the analytical force-free hard-sphere (FFHS) framework. The FFHS model assumes that the motion of the tethered spin label is slow relative to the translational diffusion of surrounding water molecules. Physically, $\tau_c$ represents the average residence time of a water molecule within the interaction distance of the spin label.

$$\xi = \frac{k_\sigma}{k_\rho} = \frac{6J(\omega_e + \omega_H;\tau_c) - J(\omega_e - \omega_H;\tau_c)}{6J(\omega_e + \omega_H;\tau_c) + J(\omega_e - \omega_H;\tau_c) + 3J(\omega_H;\tau_c)}$$

where $\omega_H$ and $\omega_e$ are the proton and electron Larmor frequencies, respectively.

Using the estimated $\tau_c$, the local translational water diffusion coefficient $D_{local}$ is be calculated based on the distance of closest approach between the spin label and water protons:

$$\tau_c = \frac{d^2}{D_{local} + D_{SL}}$$

where $D_{SL}$ is the diffusion coefficient of the spin label and $d$ is the distance of closest approach. (*2*, *3*)

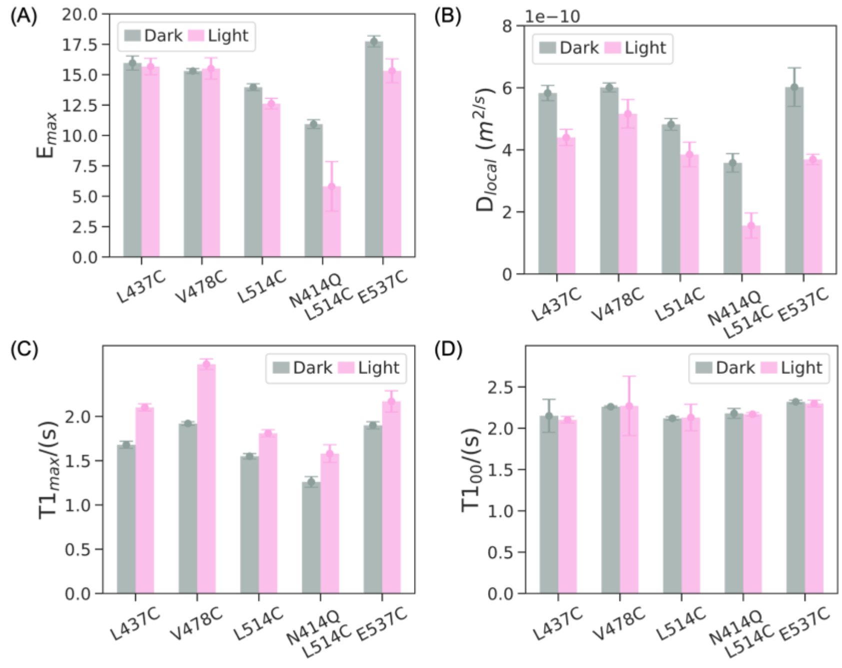

**Figure S2**. Overhauser DNP measurements at L437C, V478C, L514C, L514C(N414Q), and E537C in the dark (gray) and under blue-light activation (pink). (A) NMR signal enhancement at 2 W microwave power shows a decrease upon blue-light activation at most sites, with negligible changes at L437C and V478C. (B) Blue-light activation decreases $D_{local}$ across most sites, indicating slower local water dynamics. (C) Longitudinal relaxation times ($T1_{max}$) measured at 2 W increase in the lit state, consistent with reduced electron-driven dipolar relaxation due to decreased hydration or restricted water mobility. (D) Control measurements without spin label and without microwave irradiation show no change in $T1_{00}$ between dark and lit states, confirming that the observed effects are specific to light-induced changes detected by DNP.

### 6. Rapid-scan Time-resolved Gd-Gd EPR (TiGGER) Spectroscopy

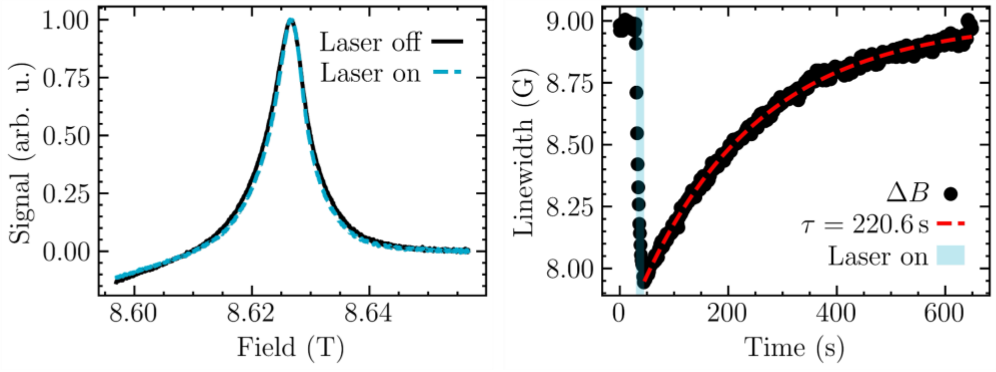

**Figure S3.** Rapid-scan time-resolved Gd-Gd EPR (*4*) measurements tracking the inter-residue distance changes between residues 406 and 537 of N414Q-mutated AsLOV2. These experiments demonstrate that at 293.2 K, N414Q AsLOV2 takes 220.6 $\pm$ 3 s to recover its dark-state configuration after a light activation event. Rapid-scan TiGGER relies on electron-electron dipolar coupling between two gadolinium spin labels at high field to elucidate inter-residue dynamics at room temperature in the solution state. Experiments were performed with the static magnetic field (B0) set at 8.62 T.

### 7. $^{17}O$ MAS NMR Spectroscopy

Purified AsLOV2 WT and N414Q samples were lyophilized for at least 48 hours and subsequently rehydrated in 40% $^{17}O$-enriched water. The rehydrated samples were packed into 3.2 mm sapphire rotors for magic angle spinning (MAS) NMR. Experiments were conducted on a Bruker AVANCE III Ultrashield Plus 800 MHz (18.8 T) narrow bore (54 mm) spectrometer, operating at a $^{17}O$ Larmor frequency of 108.44 MHz, located at the MRL Shared Experimental Facilities, University of California Santa Barbara. The MAS spinning rate was maintained at 10 kHz ± 3 Hz. For photoexcitation, a 450 nm blue laser (Laser Components USA, Inc., Bedford, NH) delivering 70 mW output was coupled into an optical fiber, transmitting approximately 15 mW to the sample space. The distal end of the fiber was mounted on the MAS probe in a manner that illuminated the sample region without direct contact with the rotor or RF coil. The $^{17}O$ 90° pulse length was calibrated to 6.0 μs using the bulk water signal from the AsLOV2 sample, which also served as a chemical shift reference at 0 ppm. Following the approach of Zhang et al.(*5*), a 180° pulse (12 μs), combined with a fixed delay $\tau_{ZC}$ = 2.75 ms (based on the measured $T_1$ of bulk water of AsLOV2), caused the bulk water signal to cross the $xy$-plane (the so-called zero-crossing), thereby minimizing its signal during detection. Setting $\tau_D$ = 0 and varying the flip angle θ enabled selective suppression of the bulk water signal, allowing for observation of wrap and bound water populations. Conversely, fixing θ to 6 μs (90° pulse) and varying $\tau_D$ permitted determination of $T_1$ values specific to the confined water species.

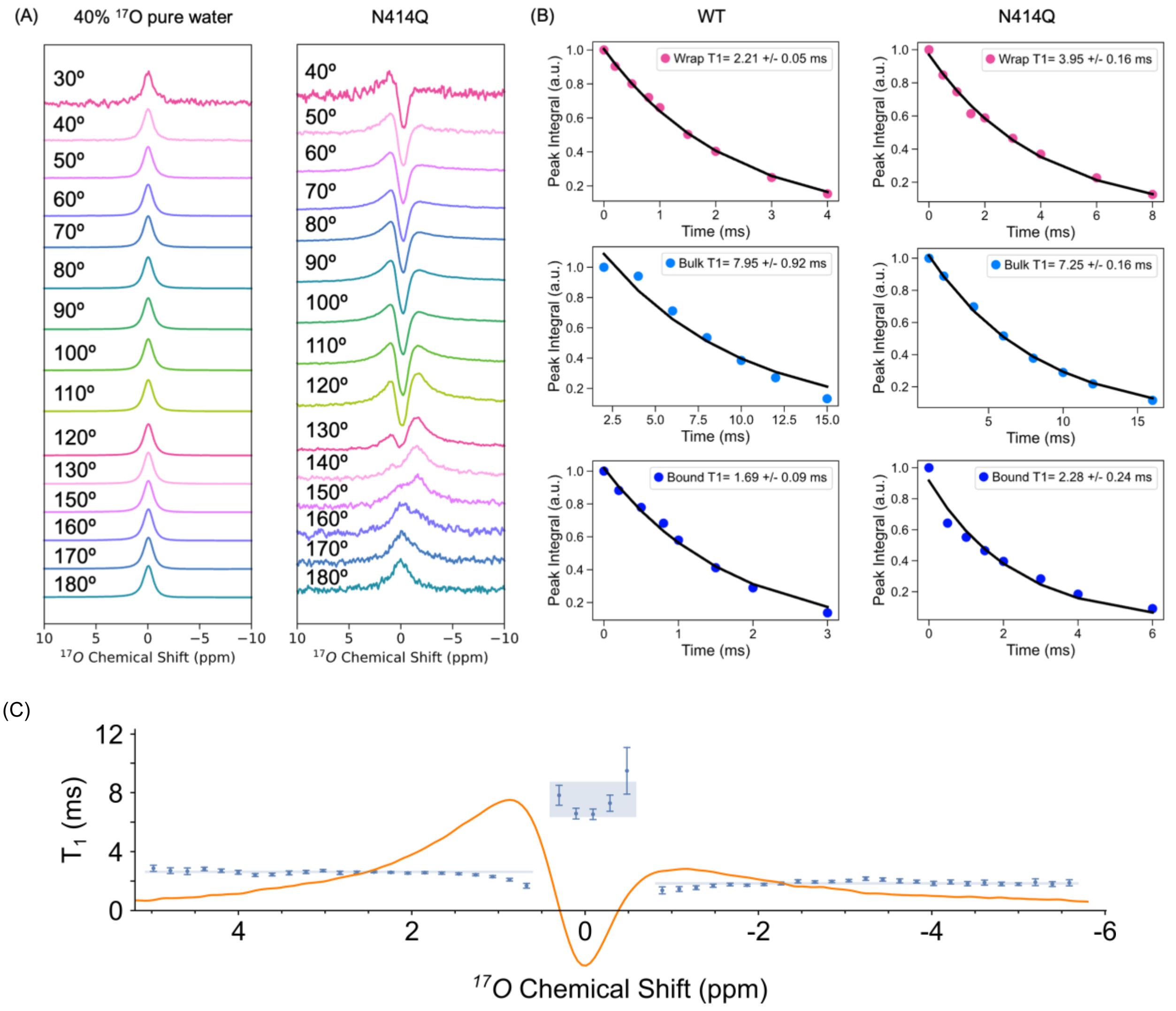

**Figure S4.** (A) Left: As a control, $^{17}O$ MAS NMR spectra of 40% $^{17}O$-enriched pure water were recorded at varying flip angles with a fixed delay $\tau_{ZC}$ = 4.89 ms (based on a measured $T_1$ = 7.05 ms of pure water). A single peak was observed across all flip angles, confirming the absence of confined water species in pure water. Right: $^{17}O$ MAS NMR spectra of AsLOV2 N414Q rehydrated in 40% $^{17}O$-enriched buffer, acquired under identical conditions except with $\tau_{ZC}$ = 3.57 ms (corresponding to a measured $T_1$ = 5.15 ms), show multiple components, indicating the presence of wrap, bulk, bound water species. (B) With the flip angle θ fixed at 6 μs (90° pulse), varying the delay $\tau_D$ enabled determination of values specific to the confined water species. Bulk water $T_1$ was 7.95 ± 0.92 ms for WT and 7.25 ± 0.16 ms for N414Q, consistent with values observed for pure $^{17}O$-enriched water. Wrap water $T_1$ was 2.21 ± 0.05 ms for WT and 3.95 ± 0.16 ms for N414Q. Bound water $T_1$ was 1.69 ± 0.09 ms for WT and 2.28 ± 0.24 ms for N414Q. (C) $T_1$ of 0.2 ppm wide slices of three regions (-6.0 to -1.0 ppm, -0.4 to 0.2 ppm and 0.9 to 11 ppm) of the spectra were calculated using a separate Mathematica script. Regions of the spectra with near zero intensities were omitted to minimize errors due to low sensitivity. The $T_1$ of each slice (with fitting uncertainty) and 99% confidence interval of the mean of slices within each region are plotted in blue, and the spectrum is overlaid in orange. T1s obtained by integral of the whole region were similar to that obtained by integral of each slice, suggesting different dynamics of spins at different chemical shift.

### 8. UV-VIS Spectroscopy

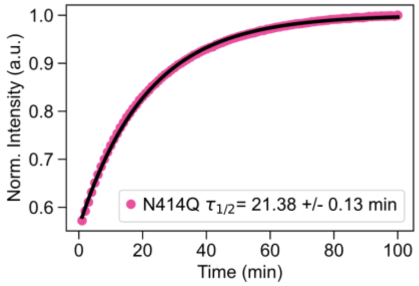

**Figure S5**. Time-resolved UV-Vis absorption of AsLOV2 N414Q monitored at 447 nm at 273 K. The lit-state recovery exhibits a lifetime decay constant of 21.38 ± 0.13 minutes, comparable to the timescale of wrap, bound, and bulk water change during dark-state recovery as observed by $^{17}O$ NMR.

### 9. Molecular Dynamics Simulations

We investigated pressure as an alternative trigger for AsLOV2 photoactivation using a two-phase molecular dynamics (MD) simulation protocol. In the first phase, we employed an implicit-solvent transfer free energy approach driven by solvent-accessible surface area (SASAR) to capture unfolding events on timescales inaccessible to conventional explicit-solvent MD (*6*), as described by Arsiccio and Shea (*7*). These simulations were conducted in AMBER (*8*) with the PLUMED SASAR plugin(*9*), running 200-ns trajectories at 298 K under pressures of 1 bar and 3 kbar. Resulting structures are shown in **Figure S9**, where we see unfolding is consistently initiated through unraveling of the A'α helix, followed by extension of the Jα helix, with high pressures accentuating Jα elongation.

In the second phase, we selected representative conformations for explicit-solvent simulations by combining RMSD-based clustering and SASAR change-point detection. RMSD clustering was performed using the GROMOS algorithm (*10*) in GROMACS (*11*) with a 0.4 nm cutoff, while change points in the SASAR time series were identified by applying a Savitzky–Golay filter (*12*) to smooth the data and using the PELT algorithm (*13*) implemented in the Python ruptures package (*14*). The former method resulted in the dominant structure shown in **Figure S6** (right), where we note clear structural changes including unraveling of the A’α helix, undocking/extension of the Jα helix, and a flattening of core β-sheets. The latter method was motivated by the theory that significant changes in water populations as a result of unfolding should be accompanied by significant changes in SASAR. These change points are shown in **Figure S8**.

For explicit-solvent simulations, we prepared selected structures along with an ambient-pressure control in GROMACS (*11*) using the CHARMM36m force field and the CHARMM-TIP3P water model (*15*). Each system was neutralized with four $Na^+$ ions and subjected to energy minimization until the maximum force fell below 1,000 kJ $mol^{-1}$ $nm^{-1}$. Equilibration was carried out in two steps: a 5-ns run at 300 K using the Berendsen thermostat and barostat (*16*), followed by a second 5-ns run at 300 K employing the velocity-rescale thermostat (*17*) and Parrinello–Rahman barostat (*18*). Production trajectories were extended for 500 ns, and analyses were confined to the 200–500 ns interval.

To analyze hydration waters we first computed protein–solvent radial distribution functions in surface mode using GROMACS rdf (*11*), identifying the second minimum at 0.38 nm as the cutoff for water inclusion. We then used MD Analysis to locate all water oxygens within this cutoff for each residue. For each central oxygen, neighboring waters within 0.35 nm were identified, and all three-body angles between the central oxygen and every pair of neighbors were computed. These angles were binned into 40 intervals spanning 40° to 180°, yielding distributions that reveal the prevalence of tetrahedral coordination(*19*, *20*). The distributions are shown for selected residues for the SASAR-determined unfolded states in water in **Figure S10**.

We quantified the tetrahedral water fraction, or wrap water, by integrating the three-body angle distribution between 100° and 120°, centered on the ideal tetrahedral angle of 109.5°. We defined bound water from the 150°–170° range of the distribution, where residence times are slowest. We quantified the icosahedral water fraction by integrating between 50° and 70°. We then correlated these values with fractional changes in residue center-of-mass distances—calculated relative to the 1 bar control over the same 200–500 ns interval—to produce the data presented in **Figure 3C**. Here we see that major changes in water population occur prior to major conformational changes of the protein itself.

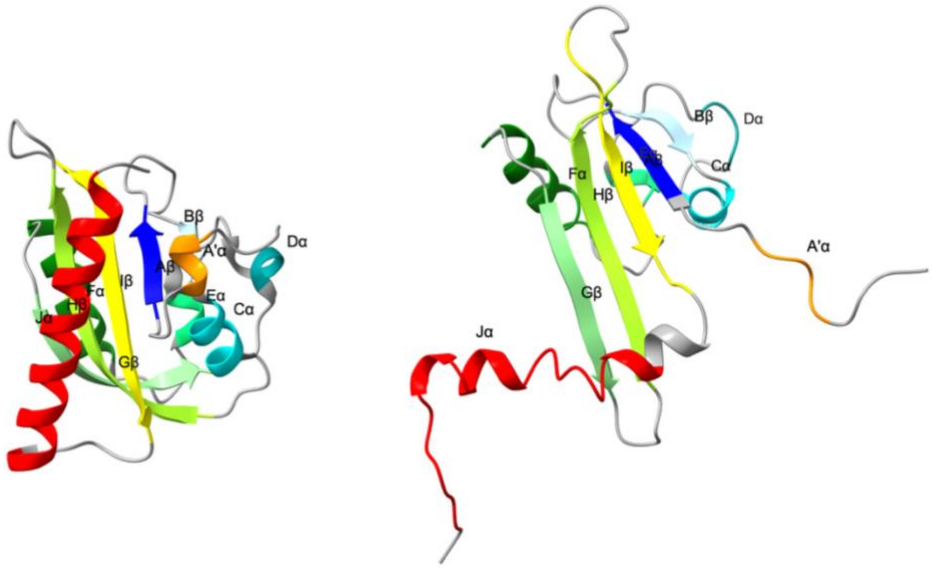

**Figure S6**. RCSB 2V1A (*21*) (residues 403–546) used for 3 kbar implicit-solvent unfolding, alongside the most populated cluster from 100–200 ns at 3 kbar, obtained by GROMOS clustering with a 0.4 nm cutoff. This dominant cluster contains roughly 11% of the sampled ensemble.

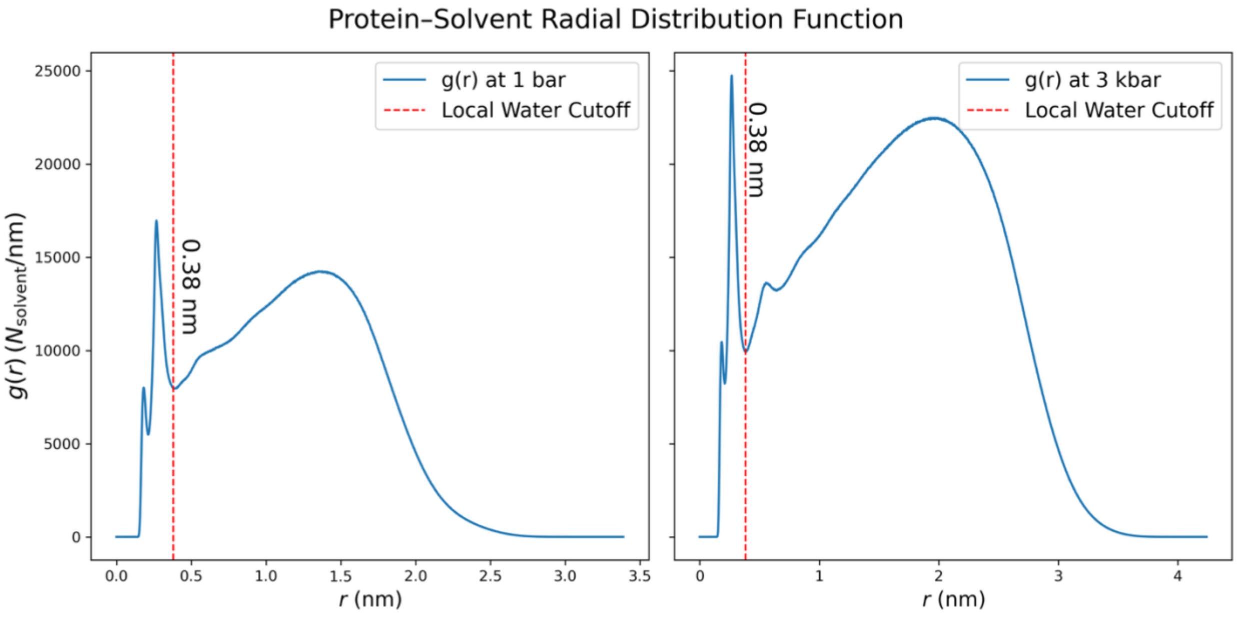

**Figure S7.** Protein–solvent radial distribution functions at 1 bar and 3 kbar, computed in surface mode with 30,001 samples. The vertical axis shows the number of solvation shell molecules per unit nm. The second minima at 0.38 nm, used for hydration analysis (*22*), are highlighted.

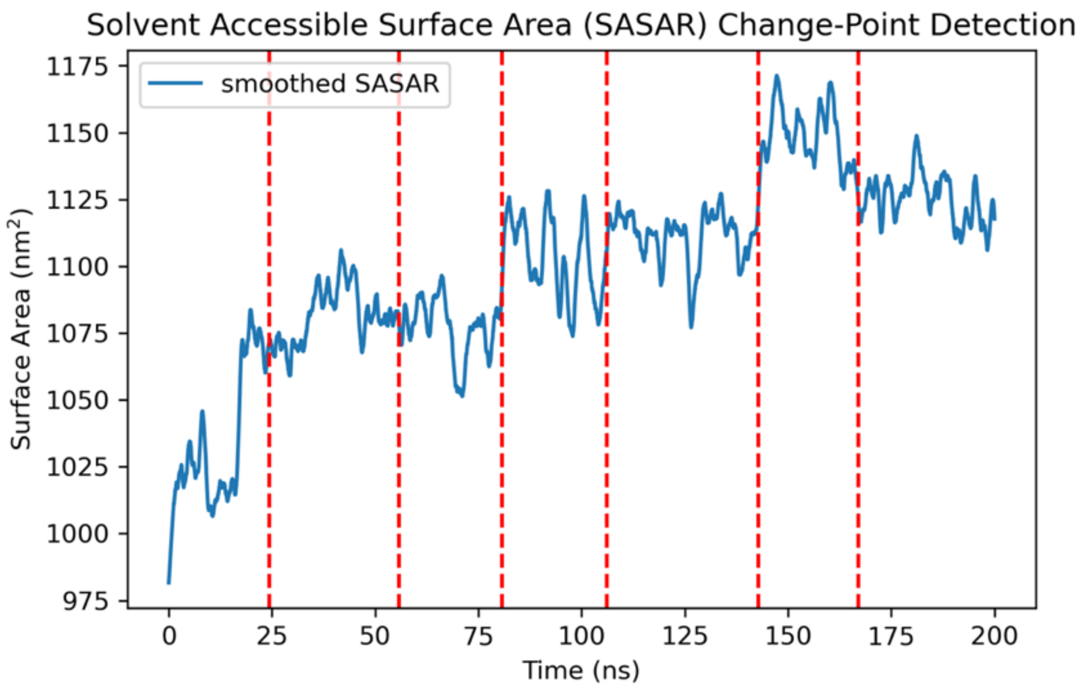

**Figure S8**. SASAR time series for the 3 kbar trajectory after smoothing with a Savitzky–Golay filter (*12*) (window size equal to 10% of the autocorrelation time, 24.316 ns). Change points at 24.320, 55.750, 80.660, 106.170, 142.750, and 167.070 ns were identified using PELT with an L2 cost; all but 80.660 ns and 167.070 ns were used for water analysis.

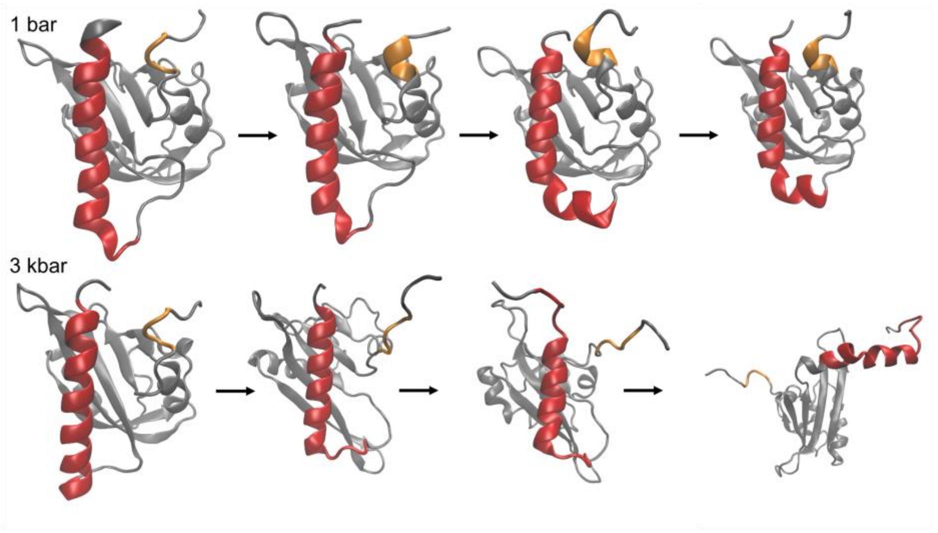

**Figure S9**. SASAR-based unfolding trajectory of AsLOV2 at 3 kbar as compared with a conventional MD trajectory in explicit solvent at ambient conditions. Snapshots taken at 0%, 25%, 50%, and 100% over each system's total simulation time. As compared with the 1 bar control, we find clear pressure-denatured states initiated by A'$\alpha$ extension, followed by J$\alpha$ unraveling until these helices point opposite to one another. Selected denatured systems (as described in the main text) were then inserted into explicit solvent for water 3BA comparison to the folded structure at ambient conditions.

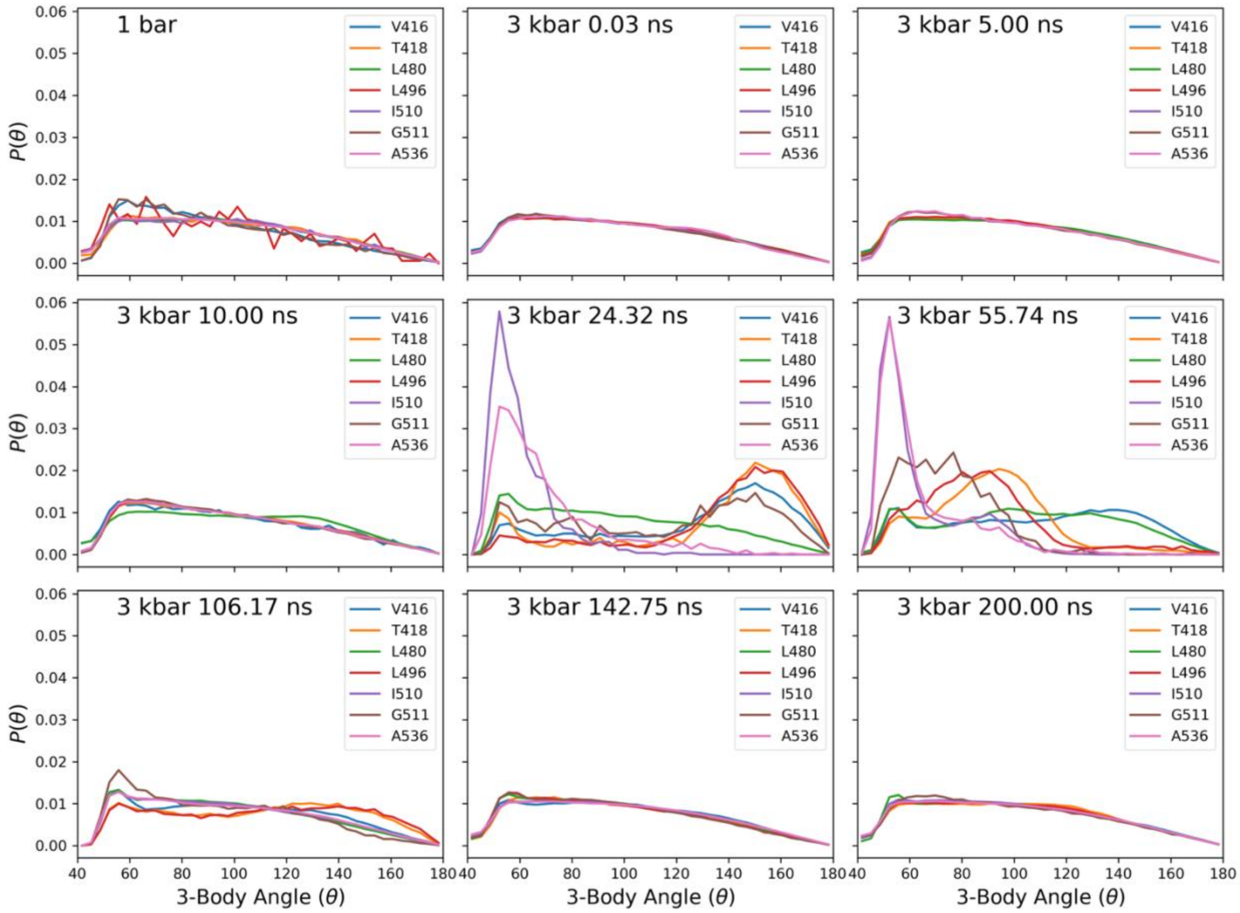

**Figure S10.** Three-body angle distributions for residues V416, T418, L480, L496, I510, G511, and A536 at 1 bar and SASAR-determined 3 kbar states in CHARMM-TIP3P water. Distributions were compiled over 200–500 ns in 40 bins spanning 40°–180°, with the 100°–120° range indicating tetrahedral coordination (*23*).

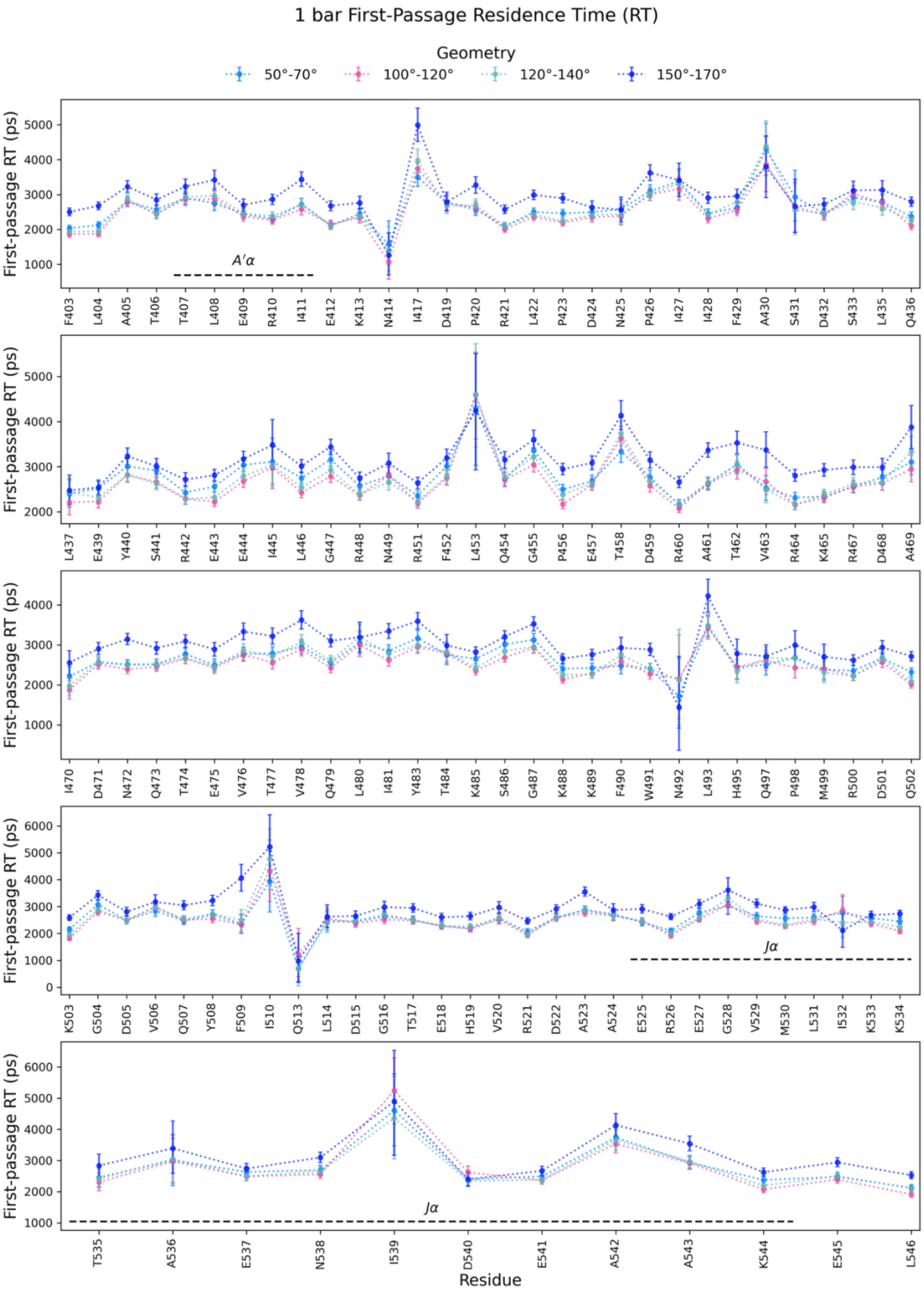

1 bar First-Passage Residence Time (RT)
Geometry
50°-70°
100°-120°
120°-140°
150°-170°
First-passage RT (ps)
A′α
F403 L404 A405 T406 T407 L408 E409 R410 I411 E412 K413 N414 I417 D419 P420 R421 L422 P423 D424 N425 P426 I427 I428 F429 A430 S431 D432 S433 L435 Q436
L437 E439 Y440 S441 R442 E443 E444 I445 L446 G447 R448 N449 R451 F452 L453 Q454 G455 P456 E457 T458 D459 R460 A461 T462 V463 R464 K465 R467 D468 A469
I470 D471 N472 Q473 T474 E475 V476 T477 V478 Q479 L480 I481 Y483 T484 K485 S486 G487 K488 K489 F490 W491 N492 L493 H495 Q497 P498 M499 R500 D501 Q502
K503 G504 D505 V506 Q507 Y508 F509 I510 Q513 L514 D515 G516 T517 E518 H519 V520 R521 D522 A523 A524 E525 R526 E527 G528 V529 M530 L531 I532 K533 K534
Jα
T535 A536 E537 N538 I539 D540 E541 A542 A543 K544 E545 L546
Jα
Residue

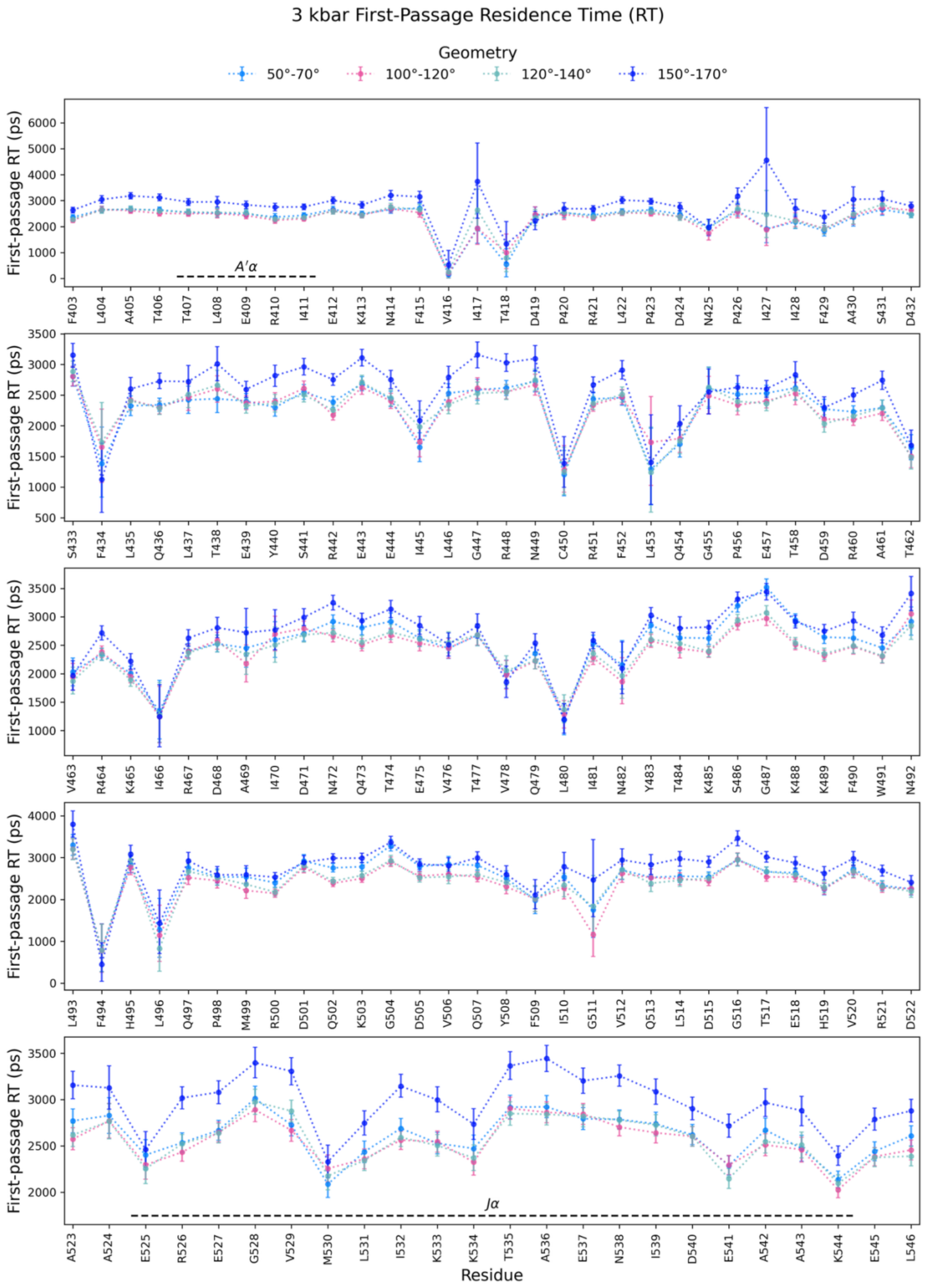
3 kbar First-Passage Residence Time (RT)
Geometry
50°-70°
100°-120°
120°-140°
150°-170°
First-passage RT (ps)
A′α
F403 L404 A405 T406 T407 L408 E409 R410 I411 E412 K413 N414 F415 V416 I417 T418 D419 P420 R421 L422 P423 D424 N425 P426 I427 I428 F429 A430 S431 D432
S433 F434 L435 Q436 L437 T438 E439 Y440 S441 R442 E443 E444 I445 L446 G447 R448 N449 C450 R451 F452 L453 Q454 G455 P456 E457 T458 D459 R460 A461 T462
V463 R464 K465 I466 R467 D468 A469 I470 D471 N472 Q473 T474 E475 V476 T477 V478 Q479 L480 I481 N482 Y483 T484 K485 S486 G487 K488 K489 F490 W491 N492
L493 F494 H495 L496 Q497 P498 M499 R500 D501 Q502 K503 G504 D505 V506 Q507 Y508 F509 I510 G511 V512 Q513 L514 D515 G516 T517 E518 H519 V520 R521 D522
Jα
A523 A524 E525 R526 E527 G528 V529 M530 L531 I532 K533 K534 T535 A536 E537 N538 I539 D540 E541 A542 A543 K544 E545 L546
Residue

**Figure S11.** Residence times of hydration water (ps) as a function of sequence position for AsLOV2 residues 403–546, calculated over the last 10 ns of production simulations at ambient and high pressure. Residence times were computed in first-passage mode, where re-entries are excluded from average lifetimes. A water is considered resident if it is within 3.8 Å of a given residue and satisfies hydrogen-bond angle criteria in one of four ranges: 50°–70°, 100°–120°, 120°–140°, or 150°–170°. To ensure statistical robustness, residues with fewer than 10 first-passage events were excluded from the analysis. Error bars represent 95% confidence intervals obtained from bootstrap resampling. Waters adopting more planar geometries (150°–170°) show markedly longer residence times than other classes, with especially pronounced differences near the flanking A'α and Jα helices that undergo large conformational changes under pressure. Historically, bound water has been defined structurally (as hydration-shell waters)(*24*), dynamically (by slow residence or reorientation)(*25*), or thermodynamically (as enthalpically stabilized)(*26*). Our results highlight a new structural–dynamic correlation: planar hydrogen-bond geometry (150°–170°) is associated with the longest residence times, suggesting a distinctive geometric motif for strongly bound water.

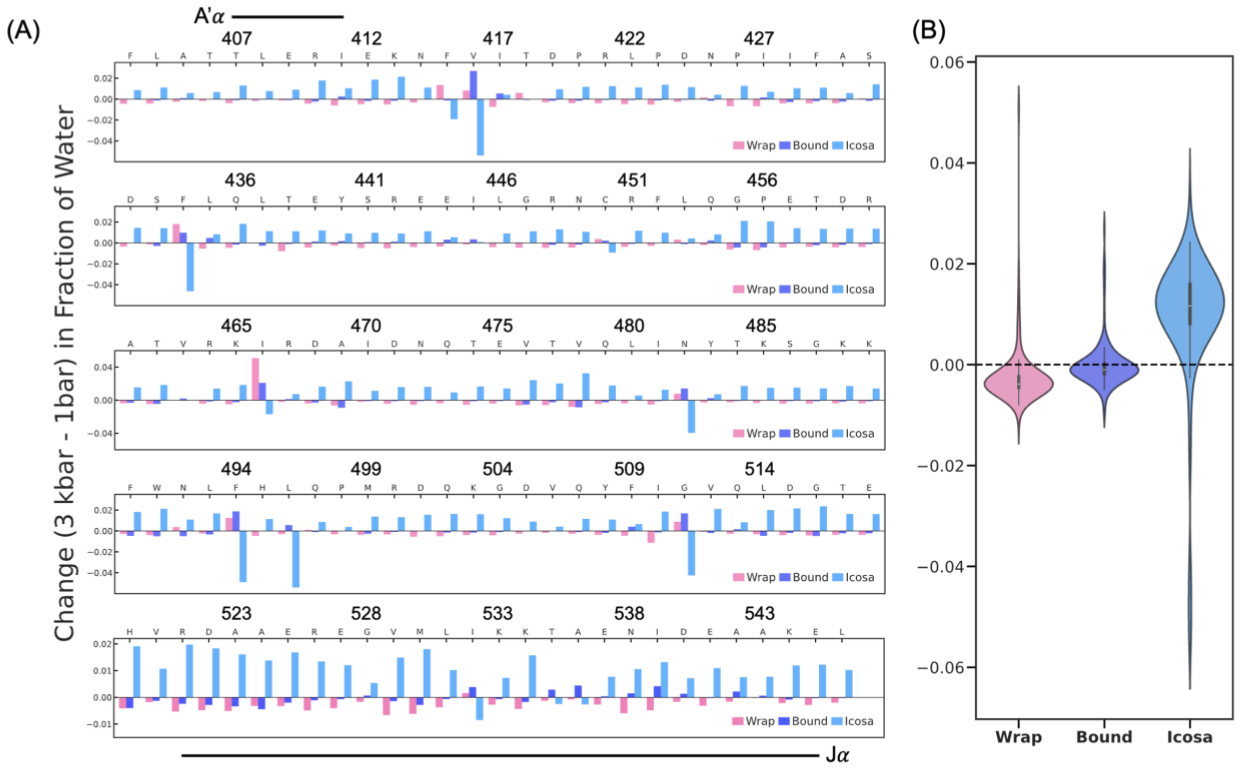

**Figure S12.** (A) Residue-wise change in the fraction of wrap (3BA: 100°–120°), bound (3BA: 150°–170°) and icosahedral (3BA: 50°–70°) water between 3 kbar and 1 bar. (B) Violin plot showing the overall pressure effect on local water populations: tetrahedral wrap water decreases, icosahedral water increases, while bound water remains nearly constant due to residue-specific behavior when pressurized from 1 bar to 3 kbar.

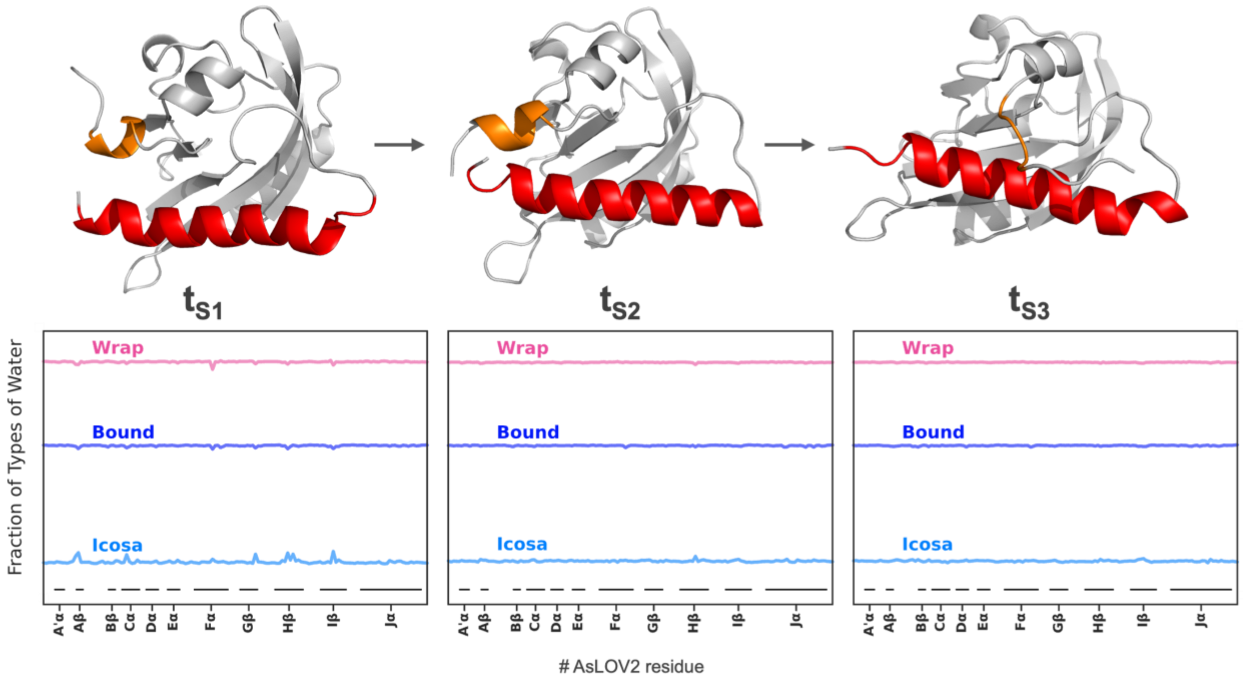

**Figure S13.** Snapshots from the 3 kbar unfolding trajectory show that wrap water remains stably associated with the protein during early unfolding events, including at $t_{S1}$ (0% of total simulation time), $t_{S2}$ (0.015% of total simulation time), and $t_{S3}$ (2.5% of total simulation time). Although the A′α helix undergoes complete unfolding within the first 2.5% of the simulation, the overall distribution of wrap, bound and icosahedral water remains largely unchanged until 12% of the total simulation time. The vertical positioning of the three plots is arbitrary.

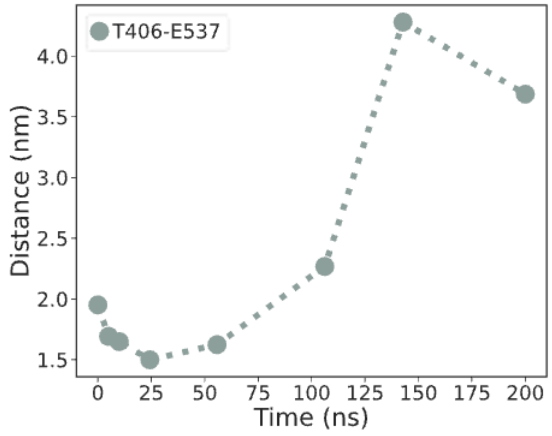

**Figure S14.** The distance between the N- and C-termini (T406 and E537) increases over time under 3 kbar pressure. By 200 ns (corresponding to 100% of the simulation time), the protein reaches a fully unfolded state with an inter-residue distance of ~4 nm. This distance is consistent with the distribution observed in high-pressure DEER measurements for light- and pressure-activated excited states.

### 10. High-Pressure Double Electron-Electron Resonance (DEER) Spectroscopy

#### 10.1. Sample Pressurization and DEER Measurements

Prior to DEER measurement, WT or N414Q AsLOV2 samples were subject to one of the following: dark conditions at atmospheric pressure; dark conditions at 3 kbar pressure; light conditions at atmospheric pressure; light conditions at 3 kbar pressure. Dark conditions refer to samples that have been prepared without exposure to blue light (λ = 470 nm), while light conditions refer to samples that have been irradiated with blue light at a wavelength of 470 nm prior to measurement. Atmospheric pressure is rated at 1 atm = 1.01325 bar.

For those samples subject to pressure, pressurization was carried out using a HUB440 high pressure generator manufactured by Pressure Biosciences Inc., designed in collaboration with the Hubbell group at UCLA (*27*). The full set up was as described in chapter 7 of (*28*) and was inspired by (*27*). To allow pressurization and subsequent DEER measurement, WT or N414Q AsLOV2 samples were loaded to 30 mm long, 1.5 m O.D., and 1.1 mm I.D. fluorinated ethylene propylene (FEP) tubes which are sealed on one end. After sample is loaded to the tube, the open end is plugged with 1.1 mm diameter silicone cord. The small outer diameter of the FEP tubes allows it to be inserted into a 15 cm long quartz tube for EPR measurement. Samples were pressurized at a magnitude of 3 kbar for 5 minutes prior to freezing under pressure using liquid nitrogen for a further 5 minutes to preserve the pressurized state. Note that 3 kbar was the maximum pressure magnitude allowed by our equipment (*28*).

Where a sample required light-irradiation, a 470 nm blue LED array light from Thorlabs Inc. (part number: LIU470A) was used. This light had an intensity of 4 mW/cm2 and was powered using a 24 V power supply. Samples, housed in a FEP tube, were subject to light irradiation from a distance of 5 cm for 60 seconds. Samples were irradiated whilst held on an ice park to slow the rate of relaxation to the dark state. Where a sample required both light-irradiation and pressure, exposure to light was carried out first, followed by immediate pressurization.

In samples that required the addition of PEG, 8 mg of PEG with a molecular weight of 20 kDa (PEG-20) was dissolved in 40 μL of WT or N414Q AsLOV2. The PEG-20 crystals were manually stirred into the AsLOV2 solution for 15 minutes under a red light in a dark room. All samples, both with and without PEG-20, were loaded into FEP tubes and plugged in the dark room under red light.

DEER measurements were carried out using a Bruker ELEXSYS E580 spectrometer at high power (150 W) Q-band frequency using an ER 5160QT-2w cylindrical resonator. A standard four-pulse DEER sequence of $(\frac{\pi}{2})_{\nu,obs} - \tau_1 - (\pi)_{\nu,obs} - (\tau_1 - t) - (\pi)_{\nu,pump} - (\tau_2 - t) - (\pi)_{\nu,obs}$ -echo was used. Here, $\nu_{obs}$ and $\nu_{pump}$ refer to pulses at the observer and pump frequencies, respectively. The timing of the pump pulse, *t*, is incremented over the course of the experiment, while the $\tau_1$ and $\tau_2$ delays are kept constant to allow a total transverse magnetisation time of $2\tau_1+2\tau_2$. To average out forbidden hyperfine transition contributions, $\tau_1$ is stepped. The echo intensity is collected over the entire time window, except for 80 ns at the beginning and end to avoid pulse-caused artefacts. The first pulse is phase-cycled. In all cases, the step repetition time (SRT) was determined as the value at which the echo amplitude was 70% of its maximum, this was 3 μs in all experiments. The other constant parameters used in all DEER experiments were $\tau_1$ = 400 ns, stepped 5 times in increments of 24 ns, $\tau_2$ = 4 μs, Δt = 8 ns, and $\nu_{pump} - \nu_{obs}$ = 80 MHz, where $\nu_{pump}$, the pump frequency, was 34.020 GHz, and $\nu_{obs}$, the observer frequency, was 33.940 GHz. All samples were measured at a temperature of 50 K. Wild type AsLOV2 samples used pump and observer π-pulse lengths of 14 ns and 22 ns, respectively, for dark conditions at atmospheric pressure and light conditions at atmospheric pressure, while lengths of 12 ns and 22 ns, respectively, were used for dark conditions at 3 kbar, and 11 ns and 22 ns, respectively, were used for light conditions at 3 kbar. Wild type AsLOV2 samples with PEG-20 used pump and observer $\pi$-pulse lengths of 16 ns and 20 ns, respectively, for dark conditions at atmospheric pressure, 12 ns and 24 ns, respectively, for light conditions at atmospheric pressure, 14 ns and 20 ns, respectively, for dark conditions at 3 kbar, and 14 ns and 22 ns, respectively, for light conditions at 3 kbar. N414Q-mutated AsLOV2 samples with and without PEG-20 used a pump $\pi$-pulse length of 14 ns and an observer $\pi$-pulse length of 22 ns for all measurements, with the exception of the light conditions at 3 kbar measurement of N414Q-mutated AsLOV2 with PEG-20 which used a pump $\pi$-pulse length of 16 ns.

For completeness, all DEER time traces were analyzed using both Gaussian modelling facilitated by DeerLab v1.1.4(*29*), and also DeerAnalysis2019 (*30*). The final populations for all AsLOV2 samples as stated in the main text were derived from the Gaussian modelling by DeerLab v1.1.4.

#### 10.2. DeerAnalysis Analysis

For analysis in DeerAnalysis, data were truncated by 600 ns to remove distortions caused by the 2+1 effect, giving a total DEER fit trace of 3296 ns in all datasets. This was not necessary in the Gaussian modelling as DeerLab is known to be minimally affected by the presence of the 2+1 effect (*31*). All backgrounds were fit using a three-dimensional homogenous distribution model, Tikhonov regularization was used to fit the corrected trace, and DeerAnalysis was allowed to determine the optimal fitting parameters. The zero time, background start value, and Tikhonov regularization parameter by the L curve criterion for each analysis are shown in Tables 1 and 2. For error analysis, the validation function in DeerAnalysis was used. The background start and end points were varied over 16 trials to determine the validation parameters. The start and end points were allowed to vary between 5% and 80% of the total length of the DEER trace.

**Table 1: DeerAnalysis2019 fitting parameters for wild type AsLOV2**

| Sample | Zero Time (ns) | Background Start (ns) | Tikhonov Parameter |
|---|---|---|---|
| 0 kbar, Dark | 332 | 968 | 1000 |
| 0 kbar, Light | 332 | 1552 | 1580 |
| 3 kbar, Dark | 333 | 936 | 501 |
| 3 kbar, Light | 332 | 400 | 1260 |
| 0 kbar, Dark, PEG-20 | 333 | 368 | 794 |
| 0 kbar, Light, PEG-20 | 332 | 344 | 1580 |
| 3 kbar, Dark, PEG-20 | 331 | 432 | 794 |
| 3 kbar, Light, PEG-20 | 332 | 400 | 1000 |

**Table 2: DeerAnalysis2019 fitting parameters for N414Q mutant of AsLOV2**

| Sample | Zero Time (ns) | Background Start (ns) | Tikhonov Parameter |
|---|---|---|---|
| 0 kbar, Dark | 334 | 994 | 1580 |
| 0 kbar, Light | 333 | 352 | 2000 |
| 3 kbar, Dark | 334 | 1120 | 1580 |
| 3 kbar, Light | 331 | 368 | 1260 |
| 0 kbar, Dark, PEG-20 | 331 | 424 | 316 |
| 0 kbar, Light, PEG-20 | 331 | 334 | 1260 |
| 3 kbar, Dark, PEG-20 | 332 | 416 | 794 |
| 3 kbar, Light, PEG-20 | 330 | 392 | 501 |

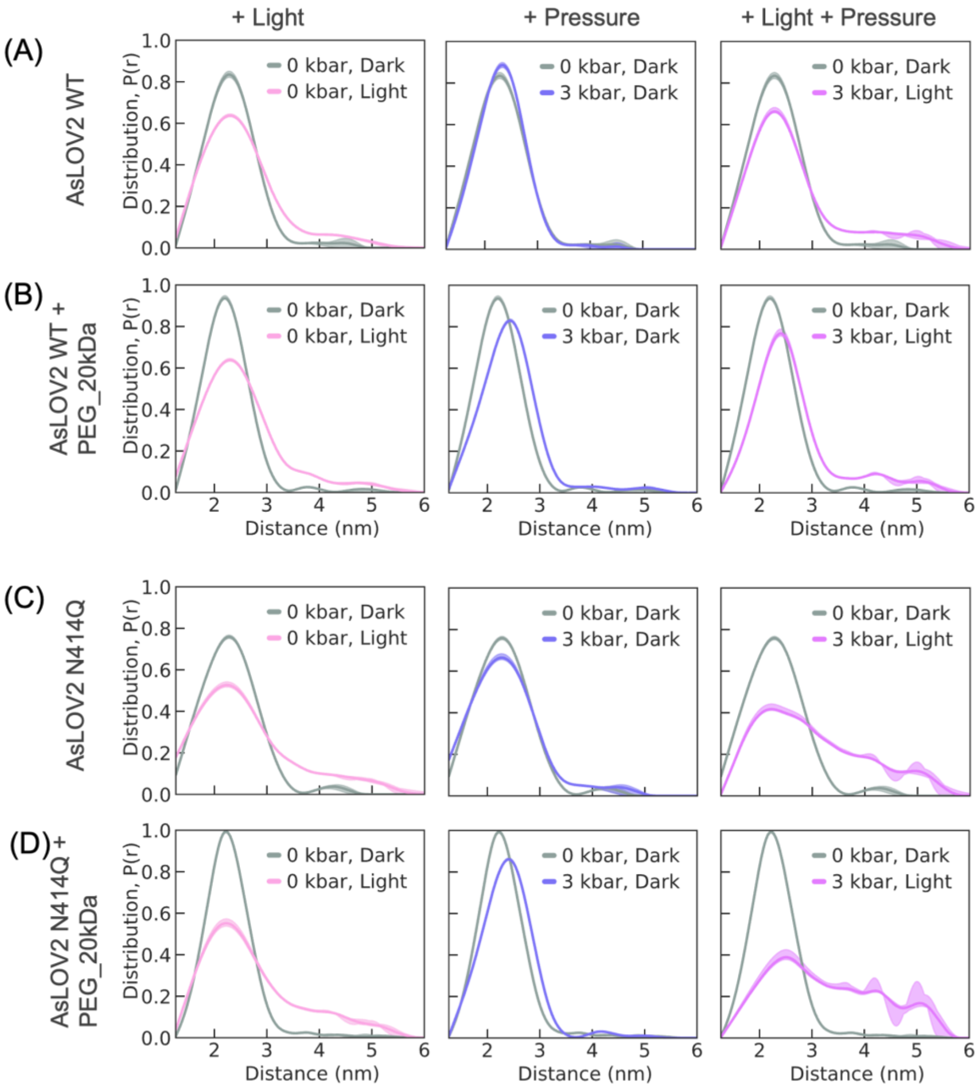

**Figure S15.** Distance distributions of AsLOV2 WT and N414Q spin-labeled at T406C and E537C with MTSL, analyzed using DeerAnalysis.

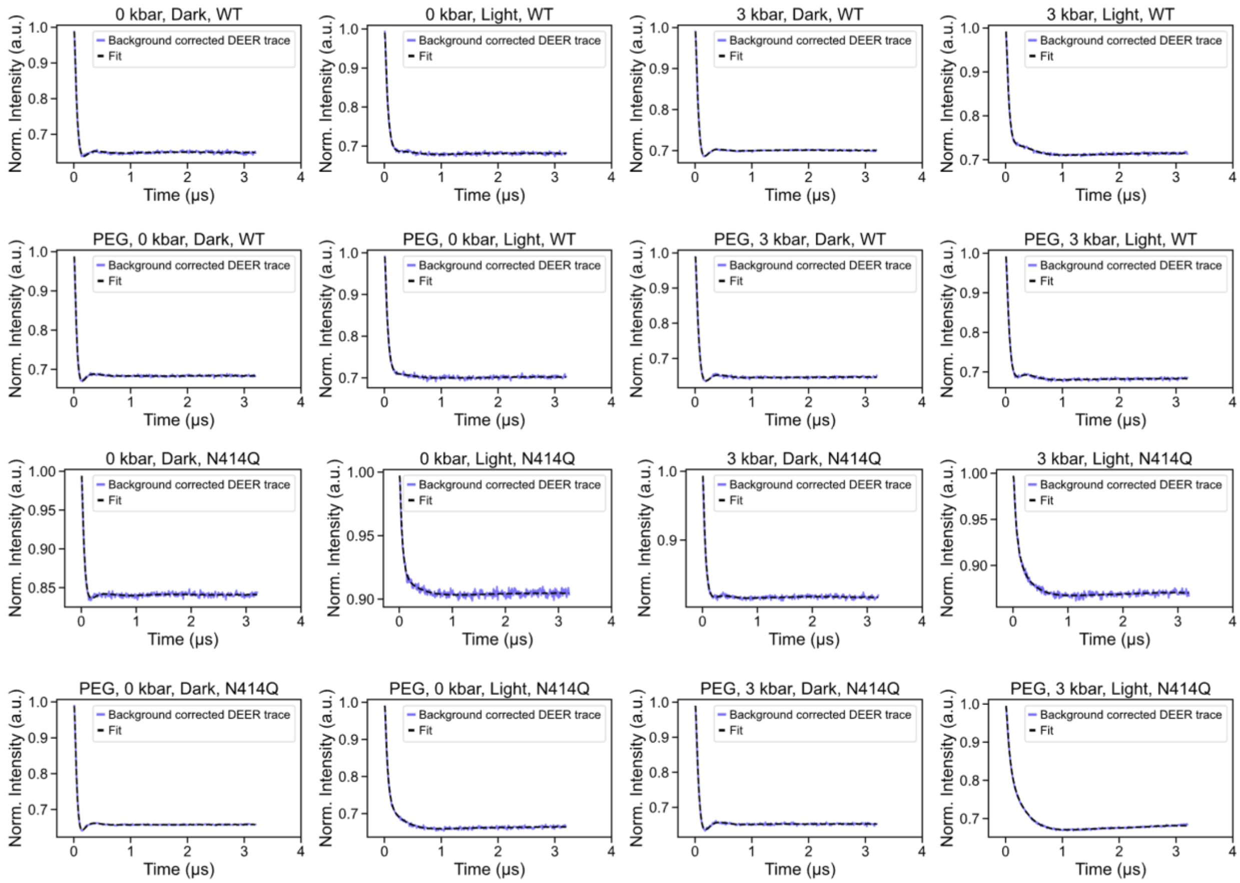

**Figure S16**. All background signals were fitted using a three-dimensional homogeneous spin distribution model. Tikhonov regularization was applied to the background-corrected DEER traces, and optimal fitting parameters were determined using DeerAnalysis.

#### 10.3. DeerLab Analysis

A model-based sum of Gaussians analysis was carried out using DeerLab v1.1.4(*29*). Data was processed in its full, original form with no truncation. A distance vector was applied between 0.5 nm and 8 nm, stepped in 0.01 nm increments. The experiment model was “ex_4pdeer” and used one pathway, while a three-dimensional homogenous background was fit using the “dd_hom3d” function. Data were fit using one or two restrained Gaussians using the functions “dd_Gauss” or “dd_Gauss2”, respectively. The first Gaussian, or only Gaussian in the case of one Gaussian fits, was restrained between 1 nm and 3 nm, while the second Gaussian, if used, was restrained between 3 nm and 6 nm. The first Gaussian restraint is based on the Multiscale Modelling of Macromolecules (MMM) software (version 2018.2)(*32*) prediction of the distance between MTSL labels at sites 406 and 537 in a crystallized structure, which was calculated to be ~2.6 nm. The second Gaussian restraint is based on the light state (no pressure, no PEG) distance mean, which was found to be ~4 nm (as shown with DeerAnalysis above). Errors were calculated using DeerLab’s default covariance matrix approach.

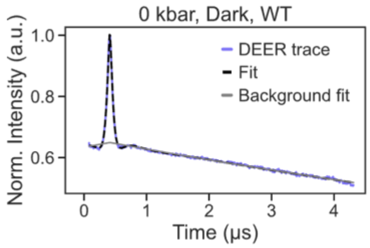

| Description | Value | 95%-Confidence interval |
|---|---|---|
| 1st Gauss Mean | 2.297 nm | (2.284, 2.309) nm |
| 1st Gauss SD | 0.442 nm | (0.428,0.457) nm |
| 2nd Gauss Mean | 4.181 nm | (4.047,4.316) nm |
| 2nd Gauss SD | 0.331 nm | (0.182,0.479) nm |
| 1st Gauss amp | 1.944 | (1.938,1.950) |
| 2nd Gauss amp | 0.066 | (0.061,0.072) |

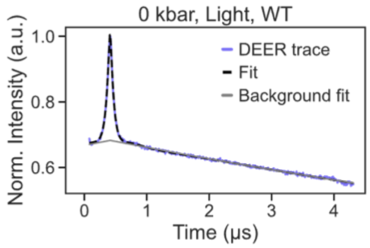

| Description | Value | 95%-Confidence interval |
|---|---|---|
| 1st Gauss Mean | 2.342 nm | (2.314,2.370) nm |
| 1st Gauss SD | 0.536 nm | (0.496,0.576) nm |
| 2nd Gauss Mean | 4.097 nm | (3.742,4.452) nm |
| 2nd Gauss SD | 0.670 nm | (0.437,0.904) nm |
| 1st Gauss amp | 1.809 | (1.801,1.817) |
| 2nd Gauss amp | 0.209 | (0.201,0.217) |

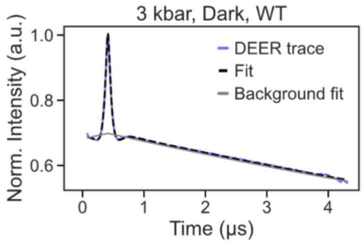

| Description | Value | 95%-Confidence interval |
|---|---|---|
| Gauss Mean | 2.323 nm | (2.315,2.331) nm |
| Gauss SD | 0.452 nm | (0.445,0.460) nm |

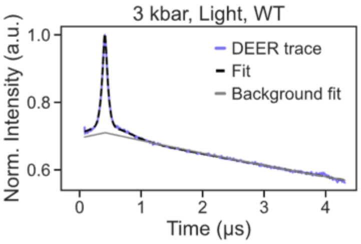

| Description | Value | 95%-Confidence interval |
|---|---|---|
| 1st Gauss Mean | 2.310 nm | (2.293,2.327) nm |
| 1st Gauss SD | 0.463 nm | (0.434,0.492) nm |
| 2nd Gauss Mean | 4.012 nm | (3.658,4.365) nm |
| 2nd Gauss SD | 1.097 nm | (0.850,1.343) nm |
| 1st Gauss amp | 1.572 | (1.564,1.581) |
| 2nd Gauss amp | 0.436 | (0.427,0.444) |

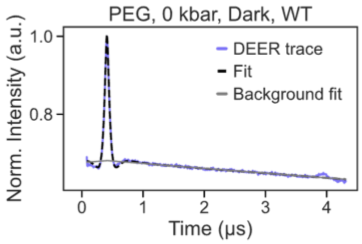

| Description | Value | 95%-Confidence interval |
|---|---|---|
| 1st Gauss Mean | 2.232 nm | (2.220,2.244) nm |
| 1st Gauss SD | 0.409 nm | (0.388,0.430) nm |
| 2nd Gauss Mean | 3.982 nm | (3.000,7.580) nm |
| 2nd Gauss SD | 1.310 nm | (0.050,2.500) nm |
| 1st Gauss amp | 1.955 | (1.945,1.965) |
| 2nd Gauss amp | 0.051 | (0.041,0.061) |

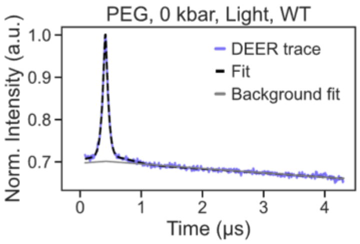

| Description | Value | 95%-Confidence interval |
|---|---|---|
| 1st Gauss Mean | 2.284 nm | (2.255,2.312) nm |
| 1st Gauss SD | 0.482 nm | (0.416,0.548) nm |
| 2nd Gauss Mean | 3.297 nm | (3.000,4.362) nm |
| 2nd Gauss SD | 1.267 nm | (0.831,1.703) nm |
| 1st Gauss amp | 1.483 | (1.468,1.498) |
| 2nd Gauss amp | 0.522 | (0.507,0.537) |

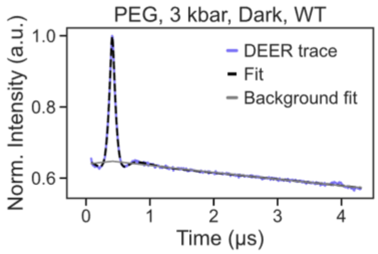

| Description | Value | 95%-Confidence interval |
|---|---|---|
| 1st Gauss Mean | 2.401 nm | (2.392,2.411) nm |
| 1st Gauss SD | 0.442 nm | (0.428,0.456) nm |
| 2nd Gauss Mean | 4.631 nm | (4.374,4.888) nm |
| 2nd Gauss SD | 0.678 nm | (0.380,0.975) nm |
| 1st Gauss amp | 1.923 | (1.918,1.928) |
| 2nd Gauss amp | 0.075 | (0.070,0.080) |

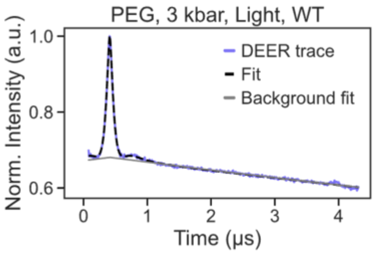

| Description | Value | 95%-Confidence interval |
|---|---|---|
| 1st Gauss Mean | 2.409 nm | (2.393,2.425) nm |
| 1st Gauss SD | 0.402 nm | (0.376,0.428) nm |
| 2nd Gauss Mean | 3.116 nm | (3.000,3.530) nm |
| 2nd Gauss SD | 1.538 nm | (1.342,1.734) nm |
| 1st Gauss amp | 1.467 | (1.455,1.479) |
| 2nd Gauss amp | 0.519 | (0.506,0.531) |

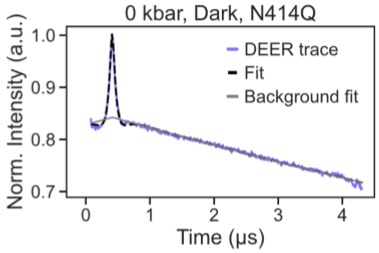

| Description | Value | 95%-Confidence interval |
|---|---|---|
| 1st Gauss Mean | 2.328 nm | (2.311,2.345) nm |
| 1st Gauss SD | 0.480 nm | (0.456,0.504) nm |
| 2nd Gauss Mean | 4.574 nm | (4.574,4.574) nm |
| 2nd Gauss SD | 2.433 nm | (2.433,2.433) nm |
| 1st Gauss amp | 1.880 | (1.841,1.920) |
| 2nd Gauss amp | 1.84e-12 | (0.00e+00,0.044) |

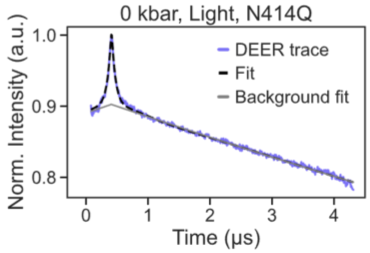

| Description | Value | 95%-Confidence interval |
|---|---|---|
| 1st Gauss Mean | 2.376 nm | (2.312,2.441) nm |
| 1st Gauss SD | 0.669 nm | (0.571,0.766) nm |
| 2nd Gauss Mean | 4.598 nm | (4.275,4.921) nm |
| 2nd Gauss SD | 0.513 nm | (0.188,0.838) nm |
| 1st Gauss amp | 1.823 | (1.800,1.846) |
| 2nd Gauss amp | 0.184 | (0.162,0.207) |

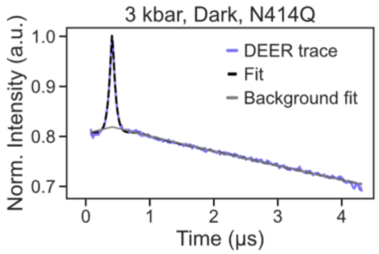

| Description | Value | 95%-Confidence interval |
|---|---|---|
| 1st Gauss Mean | 2.325 nm | (2.308,2.341) nm |
| 1st Gauss SD | 0.547 nm | (0.521,0.573) nm |
| 2nd Gauss Mean | 4.316 nm | (4.068,4.564) nm |
| 2nd Gauss SD | 0.376 nm | (0.118,0.634) nm |
| 1st Gauss amp | 1.945 | (1.934,1.955) |
| 2nd Gauss amp | 0.065 | (0.055,0.076) |

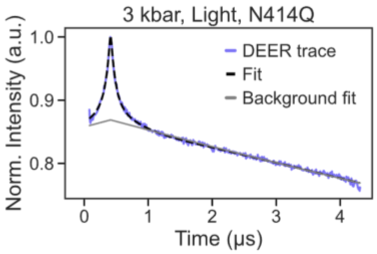

| Description | Value | 95%-Confidence interval |
|---|---|---|
| 1st Gauss Mean | 2.489 nm | (2.305,2.672) nm |
| 1st Gauss SD | 0.706 nm | (0.548,0.863) nm |
| 2nd Gauss Mean | 4.350 nm | (3.660,5.039) nm |
| 2nd Gauss SD | 0.825 nm | (0.431,1.219) nm |
| 1st Gauss amp | 1.476 | (1.457,1.494) |
| 2nd Gauss amp | 0.533 | (0.515,0.551) |

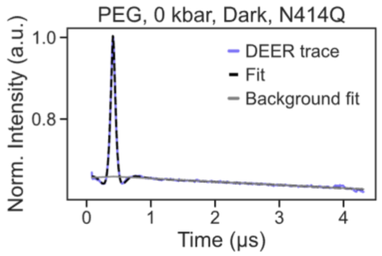

| Description | Value | 95%-Confidence interval |
|---|---|---|
| 1st Gauss Mean | 2.235 nm | (2.228,2.242) nm |
| 1st Gauss SD | 0.395 nm | (0.381,0.408) nm |
| 2nd Gauss Mean | 3.505 nm | (3.000,5.160) nm |
| 2nd Gauss SD | 1.031 nm | (0.186,1.876) nm |
| 1st Gauss amp | 1.929 | (1.924,1.934) |
| 2nd Gauss amp | 0.082 | (0.077,0.087) |

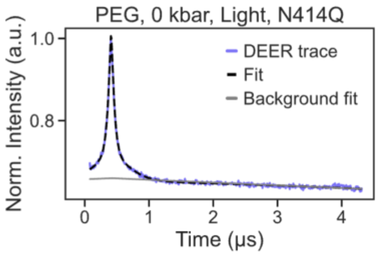

| Description | Value | 95%-Confidence interval |
|---|---|---|
| 1st Gauss Mean | 2.241 nm | (2.216,2.265) nm |
| 1st Gauss SD | 0.460 nm | (0.395,0.525) nm |
| 2nd Gauss Mean | 3.324 nm | (3.000,3.877) nm |
| 2nd Gauss SD | 1.274 nm | (1.027,1.521) nm |
| 1st Gauss amp | 1.083 | (1.071,1.096) |
| 2nd Gauss amp | 0.935 | (0.922,0.947) |

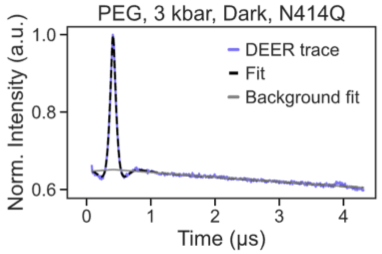

| Description | Value | 95%-Confidence interval |
|---|---|---|
| 1st Gauss Mean | 2.378 nm | (2.365,2.392) nm |
| 1st Gauss SD | 0.412 nm | (0.393,0.432) nm |
| 2nd Gauss Mean | 3.000 nm | (3.000,4.882) nm |
| 2nd Gauss SD | 1.527 nm | (0.691,2.363) nm |
| 1st Gauss amp | 1.824 | (1.812,1.837) |
| 2nd Gauss amp | 0.130 | (0.118,0.143) |

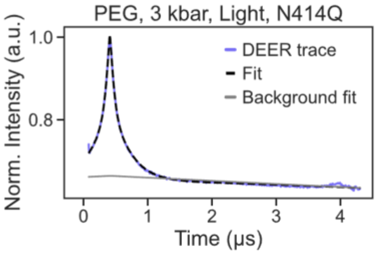

| Description | Value | 95%-Confidence interval |
|---|---|---|
| 1st Gauss Mean | 2.414 nm | (2.379,2.450) nm |
| 1st Gauss SD | 0.381 nm | (0.285,0.477) nm |
| 2nd Gauss Mean | 3.510 nm | (3.284,3.736) nm |
| 2nd Gauss SD | 1.419 nm | (1.301,1.536) nm |
| 1st Gauss amp | 0.421 | (0.414,0.427) |
| 2nd Gauss amp | 1.605 | (1.599,1.612) |

**Figure S17.** Gaussian analysis using DEERLab.

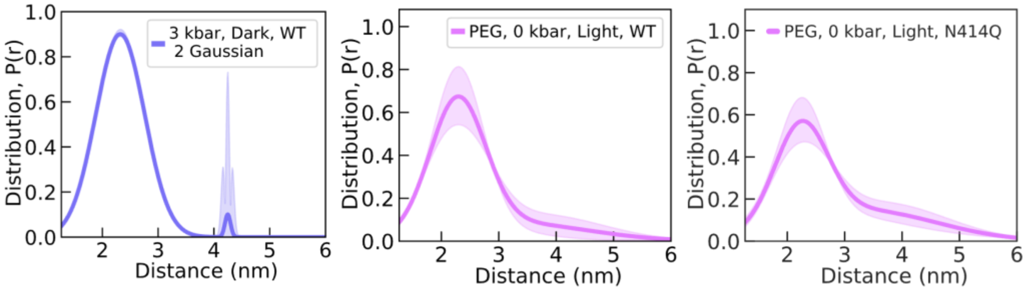

**Fig. S18.** Left: Distance distribution for AsLOV2 WT at 3 kbar in the dark, fitted with a two-Gaussian model, shows an anomalous peak at ~4 nm; therefore, this dataset was ultimately analyzed using a single-Gaussian fit, as presented in the main text (Fig. 4). Middle: Two-Gaussian fit of distance distribution for AsLOV2 WT at 0 kbar under blue-light activation in the presence of PEG (not shown in main text). Right: Two-Gaussian fit of distance distribution for AsLOV2 N414Q at 0 kbar under blue-light activation in the presence of PEG (not shown in main text).

### 11. High-Pressure three-dimensional NMR Spectroscopy

Solution NMR experiments were performed on a Bruker Avance III HD 700 MHz NMR spectrometer equipped with a 5 mm QCI-F cryoprobe, using a sample consisting of 420 μM protein in 20 mM Tris (pH 7.5), 150 mM NaCl, 6mM NaN3, 20% D2O. Samples were loaded into a pressure-resistant NMR cell (Daedalus Innovations), and pressurized to various levels between 20 and 2500 bar applied through a Xtreme-60 Syringe Pump from the same vendor. The sample was equilibrated at 298.2 K, 20 bar before confirming integrity of protein with 1D $^{1}$H and 2D $^{15}$N/$^{1}$H HSQC NMR spectra. 3D HNCO experiments (32 scans, 2048 x 60 x 100 complex points) were collected using 12% non-uniform sampling (NUS)(*33*) at 20, 500, 750, 1000, 1250, 20, 1750, 2000, 20, 2500, and 20 bar, taking 5 min at each new pressure to equilibrate starting a 7 h 43 min data acquisition. All data from 20 bar experiments were used to check for protein stability and reversibility of any pressure-induced conformational changes. All NMR spectra were processed using NMRpipe(*34*) and analyzed using NMRFx Analyst(*35*).

Chemical shift assignments were obtained from standard triple resonance HNCACB and CBCA(CO)NH experiments on $^{13}$C/$^{15}$N-enriched AsLOV2 (404-546) samples. NMR spectra for assignments were also processed using NMRpipe and analyzed with CcpNMR Analysis(*36*). Assignments were done through manual inspection of spectral connectivity patterns and confirmed by comparing to previously published NMR chemical shift assignments of AsLOV2(404-560, BMRB 26854)(*37*).

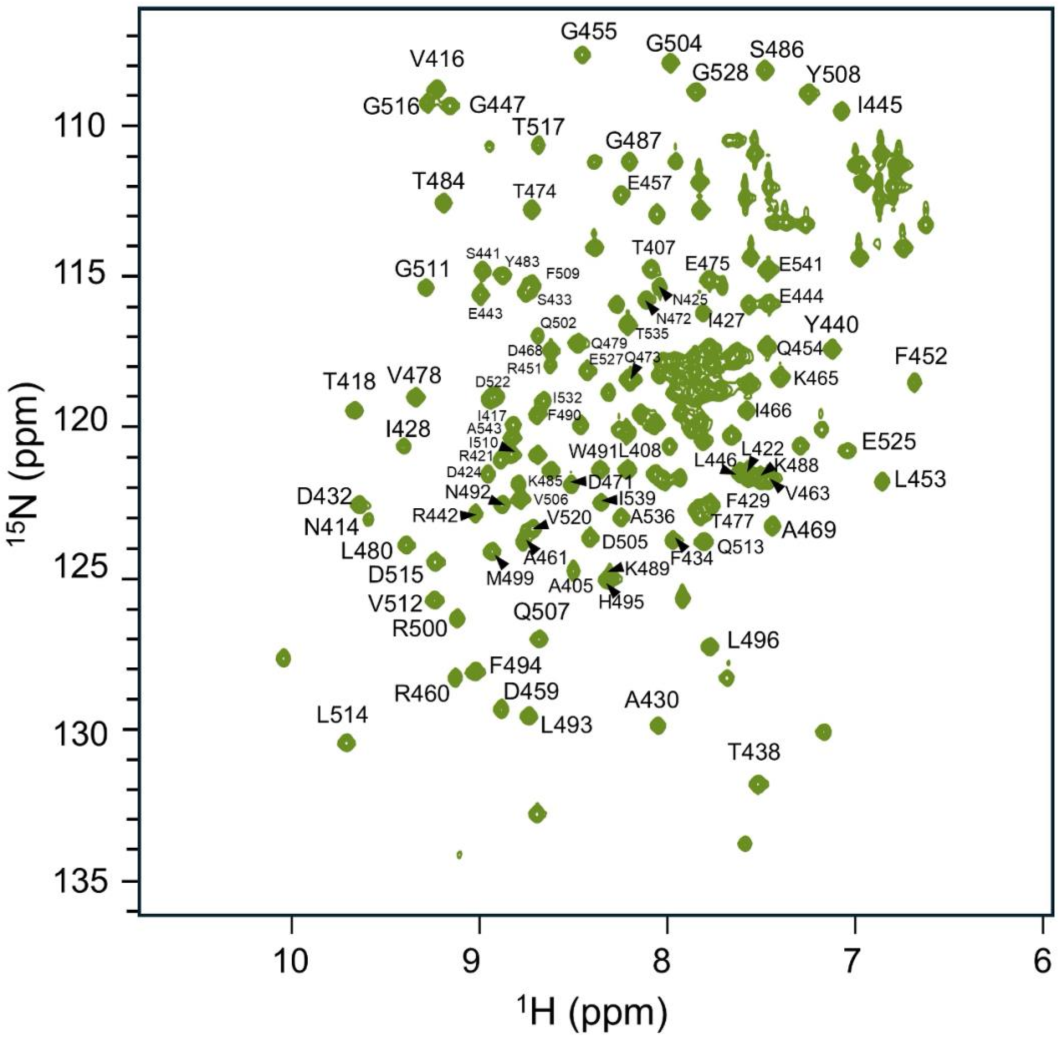

**Figure S19**: 2D $^{1}H/^{15}N$ HSQC spectrum of uniformly-$^{13}C/^{15}N$-labeled AsLOV2 (404-546) recorded at 700 MHz and 298.2 K. Backbone amide resonances are labeled with the corresponding assignments.

For the pressure jump HNCO series, peak locations were analyzed at different pressures, allowing assignments to be transferred from 1 bar reference numbers to any of the higher pressures where data were collected. Pressure-dependent changes in chemical shifts were calculated and fitted using a customized version of NMRFx(*35*, *38*, *39*):

$$\delta_i = a_i + b_i p + c_i p^2 \qquad (1)$$

where $p$ is the pressure (bar) and $b_i$ (parts per million per bar) and $c_i$ (parts per million per square bar) are the first- and second-order pressure dependent coefficients on chemical shifts for the $i$th residue respectively.

To aggregate the information of three NMR-active nuclei at any given peptide bond, we generated a scaled version of nonlinear coefficient, $c_i$, as indicated below:

$$|c_i|_{scaled} = \left(\frac{|c_i|_H}{|c_i|_{H,max}}\right) + \left(\frac{|c_i|_N}{|c_i|_{N,max}}\right) + \left(\frac{|c_i|_{C_{(i-1)}}}{|c_i|_{C_{(i-1)},max}}\right) \qquad (2)$$

Residue specific changes in chemical shift at increasing pressure was calculated by subtracting the chemical shift at 20 bar from the chemical shift at all subsequent pressures:

$$\Delta\delta\ ppm = p_x - p_{20} \quad (3)$$

The changes in chemical shifts of residue assignments for a particular nuclei were plotted using ggplot2 in RStudio(*40*).

The pressure-dependent chemical shift changes for $^{15}N$ show a general trend of downfield shifts for each residue with some deviations in the J$\alpha$ and A'$\alpha$ helical regions and an average shift of 0.421 $\pm$ 0.474 ppm at 2500 bar. This result aligns with what is typically observed for nitrogen and can be caused by the shortening of hydrogen bond distances as well as changes in the backbone dihedral angles ($\phi$ and $\varphi$)(*41*). The $^{1}H$ plots display a more dynamic range of outcomes with both upfield and downfield chemical shift changes observed and an average shift of 0.0356 $\pm$ 0.0936 ppm at 2500 bar. Normally proton showcases more downfield shifts similar to nitrogen(*42*), however, upfield shifts have been reported in specific regions of proteins consistent with what is being observed in **Figure S20** for AsLOV2(*43*). Not much work has been done probing into the effects of high pressure on the carbonyl backbone of proteins, but the results from $^{13}CO$ are similar to what we are seeing from the proton data with an average chemical shift change of 0.0203 $\pm$ 0.138 ppm at 2500 bar. In addition, for the carbonyl data there appears to be a sequence dependent pattern relating to helical periodicity evident in the c-terminus of the J$\alpha$ helix region with all residues showcasing upfield chemical shift changes.

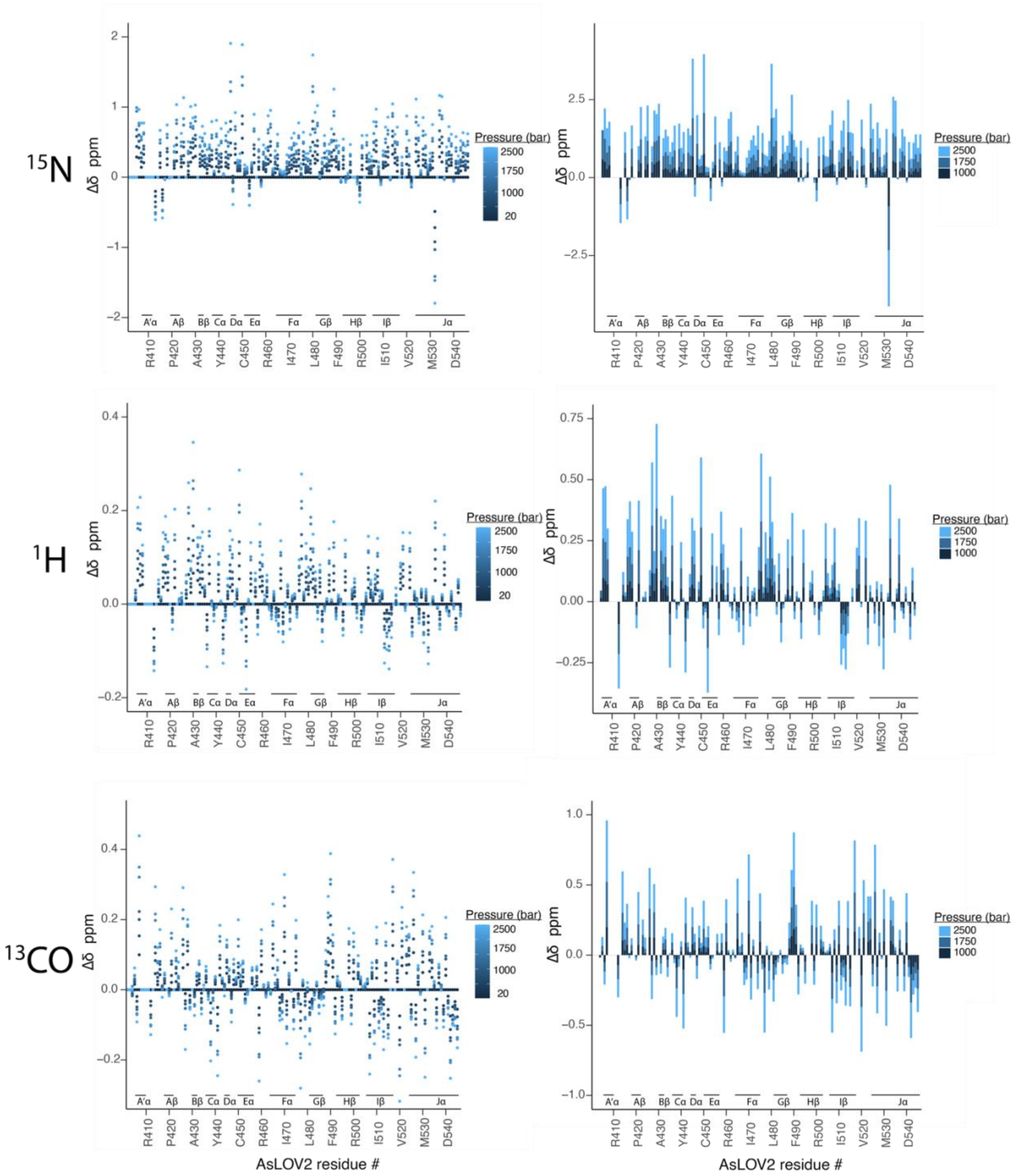

**Figure S20**. Pressure dependent chemical shift changes as measured by HNCO NMR experiments. Chemical shift differences were calculated by subtracting the chemical shift at 20 bar (baseline) from each subsequent pressure 500, 750, 1000, 1250, 1750, 2000, 2500 bar and plotting for all residue assignments as either a scatter plot (left) or as a stacked bar chart including the values at 1000, 1750, and 2500 bar (right). $^{15}N$ displays a bias towards more positive chemical shift changes while $^{1}H$ and $^{13}CO$ show uniform positive and negative chemical shift changes.
